# A cortical circuit for orchestrating oromanual food manipulation

**DOI:** 10.1101/2022.12.03.518964

**Authors:** Xu An, Yi Li, Katherine Matho, Hemanth Mohan, X. Hermione Xu, Ian Q. Whishaw, Adam Kepecs, Z. Josh Huang

**Affiliations:** Department of Neurobiology, Duke University Medical Center, Durham, NC 27710, USA; Cold Spring Harbor Laboratory, Cold Spring Harbor, NY 11724, USA; Department of Neuroscience, Canadian Centre for Behavioural Research, University of Lethbridge, Lethbridge, AB, T1K 3M4, Canada; Department of Neuroscience and Department of Psychiatry, Washington University in St. Louis, St. Louis, MO, USA; Department of Biomedical Engineering, Duke University Pratt School of Engineering, Durham, NC 27708, USA

## Abstract

The seamless coordination of hands and mouth—whether in humans eating corn on the cob or mice extracting sunflower seeds—represents one of evolution’s most sophisticated motor achievements. Whereas spinal and brainstem circuits implement basic forelimb and orofacial actions, whether there is a specialized cortical circuit that assembles these actions to enable skilled oromanual manipulation remains unclear. Here, we discover a cortical area and its cell-type-specific circuitry that govern oromanual food manipulation in mice. An optogenetic screen of cortical areas and projection neuron types identified a rostral forelimb-orofacial area (RFO), wherein activation of pyramidal tract (PT^Fezf2^) and intratelencephalic (IT^PlxnD1^) neurons induced concerted posture, forelimb and orofacial movements resembling eating. In a freely moving pasta-eating behavior, pharmacological RFO inactivation impaired the sitting posture, hand recruitment, and oromanual coordination in pasta eating. RFO PT^Fezf2^ and IT^PlxnD1^ activity was closely correlated with oromanual pasta manipulation and hand-assisted biting. Optogenetic inhibition revealed that PTs^Fezf2^ regulate dexterous hand and mouth movements while ITs^PlxnD1^ play a more prominent role in oromanual coordination. RFO forms the hub of an extensive network, with reciprocal connections to cortical forelimb and orofacial sensorimotor areas, as well as insular and visceral areas. Within this cortical network, RFO PTs^Fezf2^ project unilaterally to multiple subcortical, brainstem and spinal areas associated with forelimb and orofacial control, while ITs^PlxnD1^ project bilaterally to the entire network and the ventrolateral striatum, and can mediate concurrent forelimb and mouth movement in part through their striatal projection. Together, these findings uncover the cell-type-specific implementation of a cortical circuit that orchestrates oromanual manipulation, essential for skilled feeding.

## INTRODUCTION

The use of the hand to bring food to the mouth represents a fundamental behavioral adaptation spanning diverse vertebrate lineages ^1^. In Euarchontoglires, particularly rodents and primates, this oromanual coordination has evolved into remarkably sophisticated manipulation patterns that dramatically expand feeding opportunities ^1–5^. While sitting upright—a posture that liberates the forelimbs from body support—these animals orchestrate complex interactions between limbs, hands, mouth, and tongue to efficiently process food items ^2,3,5,6^. This evolutionarily advantageous behavior substantially broadens dietary options by enabling extraction and preprocessing of nutrient-rich components from various food sources, reducing constraints imposed by environmental niches ^2,4^.

Despite its ecological significance, the neural underpinnings of skilled oromanual coordination remain surprisingly understudied ^7^. While brainstem circuits capably implement foundational movements like licking, biting, chewing, grasping, and handling ^8–12^, a central question persists: does a dedicated cortical circuit exist to assemble these elemental actions into the sophisticated feeding behaviors observed in rodents and primates?

From Ferrier’s pioneering work ^13^ to Graziano’s influential studies ^14,15^, primate microstimulation experiments suggested cortical “action maps” containing zones for ethologically relevant movements, including concurrent hand-mouth movements. However, this framework remains controversial, with critics questioning whether artificially induced movements in head-fixed animals reveal true cortical organization for natural behaviors ^16–18^. While rodent studies have mapped cortical regions for isolated forelimb ^9,19–26^ and orofacial ^27–31^ movements, the representation of integrated oromanual actions remains largely unexplored. More importantly, beyond the concept of an action map, each cortical area comprises diverse neuronal cell types, such as glutamatergic projection neuron types that assemble intricate local circuitry, intracortical processing streams, and cortical output channels ^32,33^. However, previous optogenetic motor mapping studies in mice mostly examined broad neuronal populations ^24–26,31^ and fell short in identifying the underlying neural circuits. Indeed, the cell type and cortical circuit basis for controlling complex oromanual behaviors remains poorly understood, especially in freely moving animals. This gap raises our central question: does a specialized neocortical circuit orchestrate hand-mouth coordination during natural feeding behaviors, or are subcortical mechanisms sufficient for these complex behaviors?

Laboratory mice offer an ideal model to address this question, as they display sophisticated hand and mouth manipulatory behavior that enables feeding on a wide variety of food items, including shelled seeds and nutrient-rich body parts of captured insects ^5,6,34–36^, and are amenable for applying the full suite of modern tools for cell type and circuit analysis ^37,38^. To uncover the neural basis of oromanual manipulation, we combined a systematic optogenetic screen of cortical areas and projection neuron (PN) types, quantitative analysis of a natural feeding behavior, cell-type resolution input-output circuit tracing, neural recording, and functional manipulation in freely moving mice. We discovered an oromanual motor area and the cell-type-specific implementation of its circuit architecture that orchestrates oromanual food manipulation.

## RESULTS

### Optogenetic identification of a cortical area that elicits fictive eating

To pinpoint cortical regions orchestrating coordinated forelimb and orofacial movements, we conducted an optogenetic activation screen across the dorsal cortex. Previous optogenetic studies in mice typically stimulated broad and mixed neuronal populations ^24–26,31^, thus obscuring potential specialized circuits for skilled oromanual behaviors. Here, we leveraged a set of newly developed mouse knock-in driver lines ^39^ that selectively target pyramidal tract (PT), corticothalamic (CT), and intratelencephalic (IT) neurons, as well as subpopulations therein. We screened six key subpopulations—PT^Fezf2^, PT^Tcerg1l^, PT^Sema3e^, IT^PlxnD1^, IT^Tbr2-E17^, CT^Tle4^—alongside a broad PN^Emx1^ line and the previously utilized *Thy1-Tg18* line ^40^ (**Fig. 1a**). In head-fixed mice, we systematically activated 128 cortical sites (3 mm x 6 mm grid) by directing 473 nm light (50 Hz, 0.5 s) through a thinned skull, while high-speed cameras recorded forelimb and orofacial movements (**Fig. 1b-c**). Among the eight driver lines, PT^Fezf2^ and IT^PlxnD1^ stood out: optogenetic stimulation of each robustly elicited coordinated forelimb-orofacial movements (**Fig. 1d**, **Extended Data Fig. 1a**, b). In contrast, broad PN^Emx1^ activation produced widely distributed motor maps (**Extended Data Fig. 1c**, g, h). In *Thy1-Tg18* mice, the sites induced strong hand-to-mouth movements tended to induce weak mouth movements, while those induced strong mouth movements tended to induce weak forelimb movement (**Extended Data Fig. 1d**, g, h). CT^Tle4^ activation induced forelimb and orofacial movements mostly in the lateral regions that largely overlap with somatosensory areas (**Extended Data Fig. 1e**, g, h). The remaining three lines of PT^Tcerg1l^, PT^Sema3e^, and IT^Tbr2-E17^ either induced weak or negligible responses (**Extended Data Fig. 1f-h**). Based on these findings, we focused subsequent experiments on PTs^Fezf2^ and ITs^PlxnD1^.

**Figure 1.**
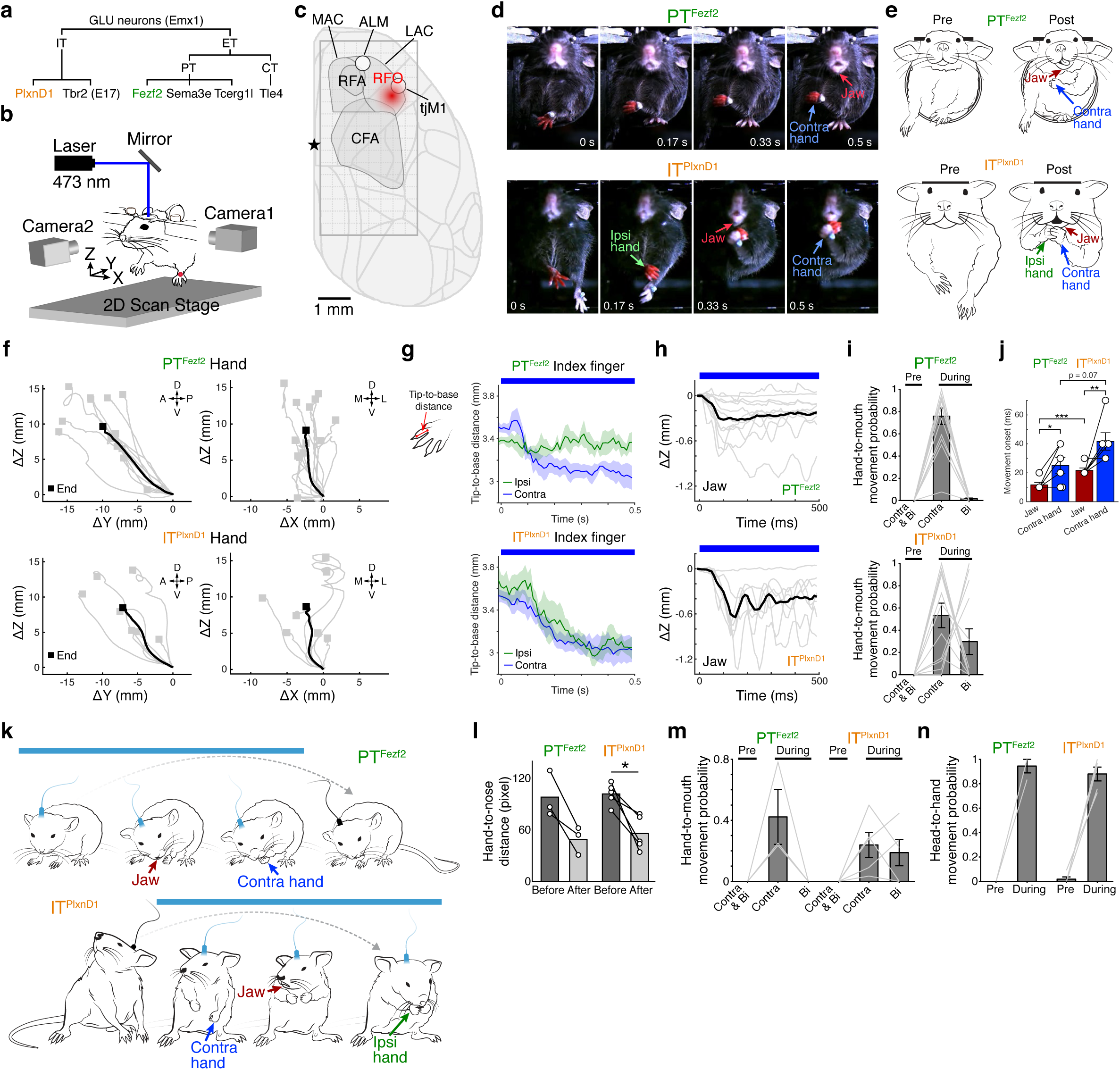
Cell-type optogenetic activation screen identifies a rostral forelimb orofacial area. **a.** PNs comprise several major classes, each comprising multiple marker-defined subpopulations used in the activation screen. IT, intratelencephalic; ET, extratelencephalic; PT, pyramidal tract; CT, corticothalamic. **b.** Schematic of optogenetic motor mapping in head-fixed mice. Nose tip denotes the coordinate origin. The red dot on the hand indicates the reflective marker for tracking. **c.** The location of RFO in relation to other motor areas. See **Extended Data Fig. 2g** for details. The rectangle grid indicates the 3mm x 6mm region mapped by optogenetic activation. Star indicates Bregma. RFA, rostral forelimb area; CFA, caudal forelimb area; MAC, medial anterior cortex; LAC, lateral anterior cortex; ALM, anterolateral motor cortex; tjM1, tongue-jaw motor cortex; RFO, rostral forelimb orofacial area. **d.** Snapshots of concurrent hand and mouth movements induced by PT^Fezf2^ and IT^PlxnD1^ activation in the RFO. See **Supplementary Videos 1, 2**. **e.** Schematic representation of forelimb and mouth movements following RFO PT^Fezf2^ and IT^PlxnD1^ stimulation. Arrows highlight movements. **f.** 2D projections of left-hand trajectories after RFO stimulation of 13 *Fezf2* mice and 7 *PlxnD1* mice. Square indicates end position. Movement directions are indicated as: A, anterior; P, posterior; D, dorsal; V, ventral; M, medial; L, lateral. **g.** The schematic depicts the measurement of the tip-to-base distance of the index finger. Right panels show changes in tip-to-base distance upon RFO PT^Fezf2^ and IT^PlxnD1^ stimulation. Bilateral and contralateral digit closure were induced by IT^PlxnD1^ and PT^Fezf2^ activation, respectively. 5 hemispheres of 3 mice for PTs^Fezf2^ or ITs^PlxnD1^. Shades around mean denote ± s.e.m. **h.** Jaw movement trajectories following RFO stimulation (11 *Fezf2* and 7 *PlxnD1* mice). **i.** Probability of observing contralateral and/or bilateral hand-to-mouth movement in a 1-s window immediately before (pre) and during RFO stimulation (13 *Fezf2* and 13 *PlxnD1* mice). **j.** Onset of jaw movement preceded that of hand movement following RFO PT^Fezf2^ or IT^PlxnD1^ activation. 6 mice for PTs^Fezf2^ or ITs^PlxnD1^. *p < 0.05, **p < 0.01, ***p < 0.005, two-way repeated measures ANOVA followed by post-hoc comparisons using the Tukey-Kramer correction. **k.** Schematic of body movements induced by RFO PT^Fezf2^ and IT^PlxnD1^ activation (blue bars) in freely moving mice. Arrows highlight movements. See **Supplementary Video 3**. **l.** Contralateral hand-to-nose distance following RFO activation in *Fezf2* (n = 3) and *PlxnD1* (n = 5) mice (*p < 0.05, two-sided paired t-test). **m, n.** Probability of observing contralateral and/or bilateral hand-to-mouth (**m**) or head-to-hand (**n**) eating-like movement in a 1-s window immediate before (pre) and during RFO stimulation (3 *Fezf2* and 5 *PlxnD1* mice). Data are mean ± s.e.m in **i, j, m**, and **n**. Movement trajectories were normalized to the start position in **f**, **h**. Black trajectories represent averages in **f**, **h**. Blue bars in **g** and **h** represent stimulation window. The mouse drawing in **b** was adapted from scidraw.io (https://scidraw.io/drawing/44).

The *Fezf2-CreER* driver line captures a majority of corticofugal neurons projecting to striatal, thalamic, collicular, brainstem, and spinal targets ^39^. Activation of PTs^Fezf2^ across the dorsal cortex of *Fezf2;Ai32* mice (expressing ChR2 in PTs^Fezf2^) revealed a topographic motor map of contralateral forelimb and orofacial movements organized along a posterior-medial to anterior-lateral axis (**Extended Data Fig. 1a**). Posterior caudal forelimb area (pCFA) stimulation induced lateral forelimb abduction with elbow extension as well as digit opening and extension (**Extended Data Fig. 2d, Supplementary Video 1**). Medial caudal forelimb area (mCFA) stimulation evoked rhythmic forelimb treading (up-down) movements (**Extended Data Fig. 2a-c, e, Supplementary Video 1**). Anterior caudal forelimb area (aCFA) stimulation induced stepping or reaching-like forelimb movements involving sequential elbow, wrist, and digit flexion followed by extension (**Extended Data Fig. 2f, Supplementary Video 1**). Notably, PT^Fezf2^ activation in an area anterolateral to the CFA induced robust and concurrent forelimb-orofacial movements, which included contralateral forelimb adduction to the body midline with hand supination and digit flexing and closing, jaw opening, and occasional tongue protrusion (**Fig. 1d-h, Supplementary Video 1**). The sequence of the forelimb and jaw movements appeared to reflect a coordinated behavior suitable for delivering food to the mouth (**Fig. 1d, e**). We named this area the Rostral Forelimb Orofacial area (RFO, **Fig. 1c**). RFO lies in the posterolateral region of the broad lateral anterior cortex (LAC, ^9^) and partially overlaps with the tongue-jaw motor cortex (tjM1), previously identified by examining orofacial movements ^41^ (**Fig. 1c, Extended Data Fig. 2g**).

The *PlxnD1-CreER* line captures a major IT population in L2/3/5A that projects bilaterally to the cortex and striatum ^39^. IT^PlxnD1^ activation in most cortical areas induced only weak or no observable forelimb movement (**Extended Data Fig. 1b**, g). Strikingly, IT^PlxnD1^ activation in the RFO generated highly coordinated bilateral forelimb-orofacial movements that resembled eating (**Fig. 1d-e, Supplementary Video 2**). These movements included jaw opening with concurrent bilateral (5/13 mice) or unilateral (8/13 mice) hand-to-mouth withdraw, flexing and closing of the digits of both hands (**Fig. 1f-i**). The bilateral forelimb movements may be attributable to the bilateral projections of ITs^PlxnD1^ in the cortex and striatum^39^. At the end of RFO IT^PlxnD1^ and PT^Fezf2^ activation, the contralateral hand was invariably moved to a consistent position close to the mouth regardless of its start position (**Extended Data Fig. 2h**), suggesting that the induced hand movement is mouth directed. Notably, IT^PlxnD1^ and PT^Fezf2^ activation induced jaw movement first, followed by a hand-to-mouth movement (**Fig. 1j**). In addition, movements induced by PT^Fezf2^ activation exhibited shorter delays compared with those induced by IT^PlxnD1^ activation (**Fig. 1j**), consistent with the fact that corticofugal projections in general have direct innervation of the brainstem and spinal motor circuitry. IT^PlxnD1^ and PT^Fezf2^ activation in a more lateral part of the RFO induced rhythmic jaw movements (instead of jaw opening) along with hand-to-mouth withdraw (**Extended Data Fig. 2i-l**). These induced forelimb and orofacial movements were robust to different stimulation frequencies and were induced primarily by long-duration stimulation (500 ms), whereas short-duration stimulation (100 ms) only induced brief and restricted movements (**Extended Data Fig. 2m**).

Because optogenetic stimulation of RFO in *Fezf2;Ai32* and *PlxnD1;Ai32* mice could also activate axons of passage of ChR2-expressing PNs from other areas, we repeated these experiments using a viral strategy to express ChR2 specifically in RFO PTs^Fezf2^ or ITs^PlxnD1^ (**Extended Data Fig. 3a**). Activating RFO PTs^Fezf2^ or ITs^PlxnD1^ was sufficient to induce synergistic forelimb and orofacial movements similar to those observed using the *Ai32* reporter mice (**Extended Data Fig. 3b-i**). Thus, our results reveal a specific cortical area (RFO), where the activation of PT or IT PNs induces forelimb-orofacial movements that resemble eating behavior.

### RFO IT^PlxnD1^ and PT^Fezf2^ activation induces fictive eating in free-moving mice

As natural feeding behavior involves the coordination of multiple body parts in addition to the forelimb and mouth, we stimulated RFO PTs^Fezf2^ and ITs^PlxnD1^ in freely moving mice. PT^Fezf2^ activation induced a shoulder adduction that raised the contralateral hand toward the body midline, with associated hand supination and digit flexion. In addition, a concurrent ipsiversive head turning and lowering brought the snout toward the contralateral hand, while the ipsilateral hand provided body support (**Fig. 1k, l, Supplementary Video 3**). Activation of RFO ITs^PlxnD1^ occasionally induced a sitting posture and concomitant bilateral shoulder adduction that brought both hands to the body midline. During the adduction, the digits flexed and closed, and both hands withdrew to the mouth (**Fig. 1k, l, Supplementary Video 3**). These results suggest that RFO PTs^Fezf2^ and especially ITs^PlxnD1^ are components of cortico-subcortical circuits mediating body posture, head, mouth, forelimb, and digit movements that are used in coordinated eating behavior. Body tracking and analysis revealed that PT^Fezf2^ and IT^PlxnD1^ activation modulated prestimulation body posture to an eating-appropriate posture. The stimulation caused an increase in body height when the initial position was low (e.g., a mouse engaged in locomotion) and a decrease in body height when the initial position was high (e.g., a mouse engaged in rearing; **Extended Data Fig. 2n**, o). Consequently, compared with head-fixed stimulation that resulted in the hand coming to the mouth, free-moving stimulation evoked a terminal eating position that could be achieved via movement of the head, hand, or both (**Fig. 1m, n**). Together, these results suggest that a defining outcome of RFO activation is to produce a functional component of eating, the juxtaposition of the mouth and hand.

### Pasta eating features oromanual coordination

To explore the role of RFO in food handling and eating, we established a “Mouse Restaurant” with a dining area that a mouse could enter to find food to eat (**Fig. 2a, Supplementary Video 4**). The restaurant was automated (including automated food delivery) so that a mouse could eat with minimal experimenter involvement (**Extended Data Fig. 4a-c**). Behavior was filmed by 3-synchronized video cameras, together with concurrent sound recording that registered the bite events associated with eating (**Fig. 2a, Extended Data Fig. 4d**). The restaurant menu consisted of a variety of food items including pellets, pasta, and seeds (see Methods). Amongst these, the angel-hair pasta deserves special mention ^42–45^. It has a consistent shape and can be tailored to a specified length (1 to 15 mm). When pieces of pasta are bitten from the stem, it features a sharp audible snap. The many movements involved in pasta-eating are distinctive and involve an interplay between the hands and the mouth. The Mouse Restaurant provided thousands of trials that were documented by millions of video frames of eating. We used DeepLabCut ^46^ to label ∼14,600 images to track 10 body parts (left and right eyes, hands, ankles, nose, tongue, jaw, and tail base) and three parts of the pasta (top, center, and bottom) (**Fig. 2b**). We analyzed over 4 million video frames to identify and annotate the action motifs of eating ^47^ (**Extended Data Fig. 4e-i**). We designed an actogram, which presents overlays of the location and action of key body parts and sensorimotor events, and co-registered bite events across an entire trial in a single graph (**Fig. 2c**).

**Figure 2.**
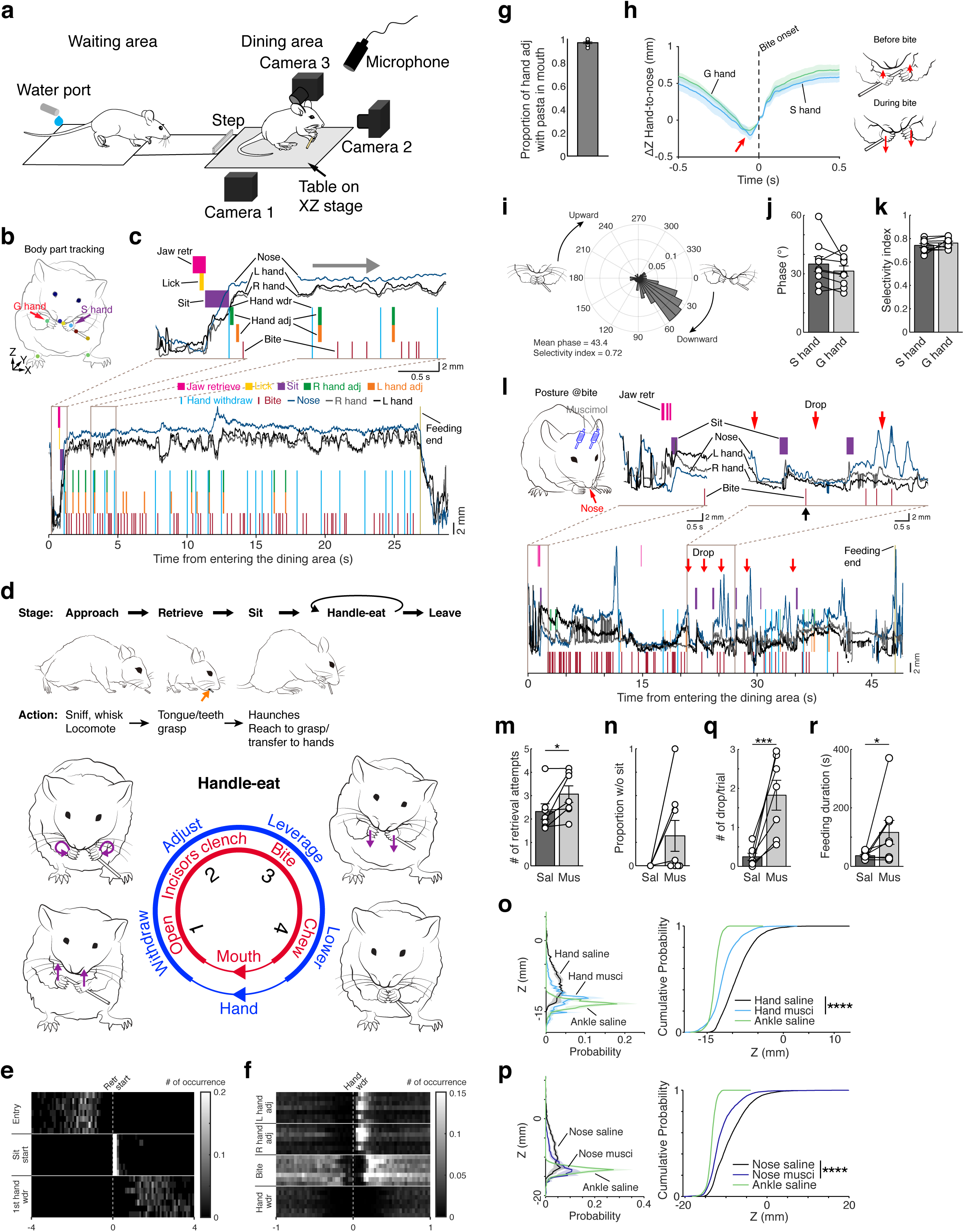
RFO is necessary for hand recruitment and oromanual coordination in pasta eating. **a.** Schematic of the Mouse Restaurant. A table mounted on an XZ stage brings food to the dining area. Three cameras record movement and a microphone records bite sound. A mouse crosses a small elevated step to enter the dining area. Also see **Extended Data Fig. 4a-c**. **b.** Pasta-eating schematic showing tracking of different body parts and pasta (colored dots). Z axis corresponds to the dorsal-ventral axis. Mice handle pasta with a support grasp (purple arrow) and a guide grasp (red arrow). **c.** Actogram of a mouse retrieving and eating a 15-mm angel-hair pasta piece. Key action motifs (color annotated) are superimposed on the Z-axis trajectories of the nose and right and left hands throughout the trial. The thick gray arrow at the top indicates the progression of a handle-eat bout. **d.** Ethogram of pasta eating, which proceeds in a sequence of stages, each consisting of multiple action motifs (top). The three schematics at the top depict actions in the approach, retrieve, and sit stages respectively. Orange arrow denotes the tongue that licks the pasta. Mice consume pasta in repeated handle-eat bouts. Bottom schematics depict a typical sequence of four coordinated hand (blue ring) and oral (red ring) actions in a handle-eat bout. Purple arrows indicate direction of hand movement. **e, f.** Heat maps showing the average number of occurrence of different actions around the retrieval start (**e**) and hand withdraw (**f**). Data from 7 mice with each row corresponding to the data of each mouse. **g.** Nearly all hand adjustments (97.0 ± 0.8 %; n = 9 mice) were made with pasta clenched in the mouth, thus involving oromanual coordination. **h.** Average hand-to-nose distance begins to increase (red arrow) before bite onset (n = 9 mice). Schematic shows hand movement immediately before and during pasta bite/snap (arrow indicates movement direction along Z axis; arrow length indicates speed). **i.** Probability distribution of the phases of left-hand movement at the time of bite from an example trial. The narrower the probability distribution the larger the selectivity index. **j**, **k**. Average hand-movement phase at the time of bite (**j**) and phase selectivity index (**k**) (n = 9 mice). **l.** Example actogram of a mouse following bilateral muscimol infusion in the RFO. Red arrows indicate pasta drops. Black arrow indicates the moment corresponding to the posture schematic in the top left panel. Note the mouse’s hunched posture with the nose close to the floor (red arrow). Also see **Supplementary Video 5**. **m, n, q, r.** RFO muscimol infusion resulted in increased retrieval attempts (**m**, n = 7 mice; *p < 0.05, two-sided paired t-test), increased number of trials in which mice consumed pasta without sitting on haunches (**n**, n = 8 mice, with one mouse never adopting a sitting posture), increased pasta drops in each trial (**q**, n = 7 mice; ***p < 0.005, two-sided paired t-test), and increased feeding duration for each pasta piece (**r**, n = 8 mice; *p < 0.05, two-sided Wilcoxon signed-rank test). **o, p.** Hand (**o**) and nose (**p**) position were lower at the time of bite following muscimol infusion compared to saline infusion as shown by the probability distribution and cumulative probability of the Z-axis position at the time of bite (n = 5 mice; ****p < 0.001, Kolmogorov-Smirnov test). Data from left ankle after saline infusion is shown as reference. Shades around mean denote ± s.e.m in **h**, **o, p**. Data are mean ± s.e.m in **g**, **j, k, m, n, q, r**. Mouse drawings in **a** were adapted from scidraw.io (https://scidraw.io/drawing/122 and https://scidraw.io/drawing/96).

The angel-hair pasta eating behavior was organized in stages, each comprising multiple characteristic action motifs involving multiple body parts (**Fig. 2c, d**). Upon entering the dining area, mice *approach* the pasta to *retrieve* it from the floor by licking (**Extended Data Figs. 4e, f, 5a**). Licking then enabled a grasp of the pasta with the incisors, which was immediately followed by a mouse adopting a *sitting posture* on the haunches (**Fig. 2d, e**). During the adoption of a sitting posture, both hands reached out to take the pasta from the teeth (**Fig. 2d**, **Extended Data Fig. 4g**). Once the pasta was held in the hands it was eaten in *handle-eat* bouts (**Fig. 2d**). After a piece of pasta was consumed, a mouse inspected the eating area by sniffing and rearing and then left the dining area.

The *handle-eat* bout was characterized by sequential hand movements that were coordinated with mouth movements to position the pasta for eating. Each bout started with a hand withdraw that brought pasta to the opening mouth (**Fig. 2c, d, Extended Data Fig. 4h**). Hand withdraw featured a specialized grasp by each hand. One hand made a *guide grasp*, which directed the end of the pasta into the mouth. The other hand made a *support grasp*, with the tips of the digits holding the pasta more distally from the mouth to feed the pasta into the mouth after each bite ^44^ (**Fig. 2b**). This advancing of the pasta into the mouth for each bite was achieved by frequent release and re-grasp movements, i.e. hand adjustments, with one or both hands (**Fig. 2c, d, Extended Data Fig. 4i**). Thus, mice used coordinated hand adjustments to position the pasta between the teeth just before the first bite of each bout (**Fig. 2c, f, Extended Data Fig. 5b**, c). Frame-by-frame analysis revealed that mice alternated holding the pasta with the mouth and hands to allow each to be sequentially repositioned as eating progressed (**Fig. 2g, Extended Data Figs. 4i, 5d**). The cooperative hand and mouth movements for pasta positioning were usually made with the pasta held at a distinct oblique angle between the hands and the teeth.

Biting pieces of pasta from the stem involved cooperative movements between the hands and the mouth. Both hands began to move rapidly downward shortly before a bite, exerting a fulcrum-like action on the pasta (**Fig. 2h, Extended Data Fig. 5e-g**). A phase analysis of the hand movement in relation to biting off a pasta piece revealed that the hand movement was tightly associated with jaw pressure exerted on the pasta that resulted in a sharp *snapping* of the pasta that produced a distinctive snap sound (**Fig. 2i-k, Extended Data Fig. 5h**, i). Pasta pieces that were snapped from the stem were then chewed (**Fig. 2d**). Thus, pasta positioning and biting all involved coordinated movements between the hands and the mouth (also see ^36^). The various movements of eating can be described as action motifs (pick-up, sit and transfer to hands, withdraw toward the mouth, hand adjustment and bite, chew). Although the details of the handle/eat movements varied from pasta to pasta, they are always recognizable and measurable (**Fig. 2d-f**).

### RFO is necessary for hand recruitment and oromanual manipulation in pasta eating

To determine whether RFO is involved in pasta eating, we suppressed neural activity by bilateral infusion of GABA_A_ receptor agonist, muscimol (**Extended Data Fig. 6a**, b). Following infusion, the mice were able to approach and locate the pasta, but they were deficient in retrieving the pasta by licking (**Fig. 2m**, **Extended Data Fig. 6c**). Mice that were able to get the pasta into their mouth often failed to adopt a sitting posture (**Fig. 2l, n, Extended Data Fig. 6d**, e). If they did get into a sitting posture, their hand use in handling the pasta was impaired. They failed to manipulate the pasta into a proper orientation for mouth grasping and biting. For example, one mouse consumed all the pasta from the floor using only its mouth (**Fig. 2n, Supplementary Video 5**). When attempting to eat in a sitting posture, the posture was hunched presumably to assist in holding the pasta but the pasta was nevertheless frequently dropped (**Fig. 2l, o-q, Extended Data Fig. 6d**, e**, Supplementary Video 5**). These oromanual impairments resulted in the mice taking significantly longer to eat (**Fig. 2r, Extended Data Fig. 6d**, e). They also resulted in the mice losing the pasta (e.g. pasta flew out of the dining area after a bite due to clumsiness of oromaunal movements) or leaving the dining area without finishing a piece of pasta. Despite these organizational impairments, the mice were not deficient in hand grip force and bite force (**Extended Data Fig. 6f**, g). Together, these results indicate that RFO regulates and coordinates multi-body-part movements, including the sitting posture, hand recruitment, and oromanual coordination, that comprise skilled hand and mouth food handling.

### RFO PN input-output connectivity forms a brain network for oromanual coordination

Having identified the RFO where PT^Fezf2^ and IT^PlxnD1^ stimulation mimics “fictive eating” we next examined its input-output wiring to understand how it might drive skilled food manipulation. Anterograde tracing revealed that PTs^Fezf2^ project prominently to multiple ipsilateral or contralateral subcortical targets in the thalamus, striatum, lateral superior colliculus (lSC), pons, and medulla (**Fig. 3a, b, Extended Data Fig. 7a-d, Supplementary Video 6**); the sparse projections to the contralateral cortical regions and striatum are likely derived from a small set of *PlxnD1*^+^ neurons within the *Fezf2*^+^ population ^39^. The brainstem targets of PTs^Fezf2^ include multiple command centers for forelimb and orofacial actions such as forelimb reaching and handling (PARN/latRM) ^8,12^, grasping (PARN, MDRN) ^8,12,48^, jaw opening and closing (SUT, PSV, SPV, IRN) ^11,49,50^, licking (PSV, SPV, IRN) ^11,49,50^, and whisking (PSV, SPV, IRN) ^11,50,51^. The PT^Fezf2^ projection crosses midline at the pyramidal decussation to innervate the spinal cord (**Extended Data Fig. 7c**). These results suggest that PTs^Fezf2^ in the RFO have direct access to both forelimb and orofacial subcortical and spinal targets. A recent study suggests that PT neurons in the LAC generally do not project to the spinal cord ^9^, seemingly inconsistent with our finding on PTs^Fezf2^ in the RFO, which resides in the posterolateral region of the LAC. Since LAC is a large territory, it is possible that previous anterograde injections and AAV-retro tropism ^52^ may have not included the RFO and missed the corticospinal neurons therein.

**Figure 3.**
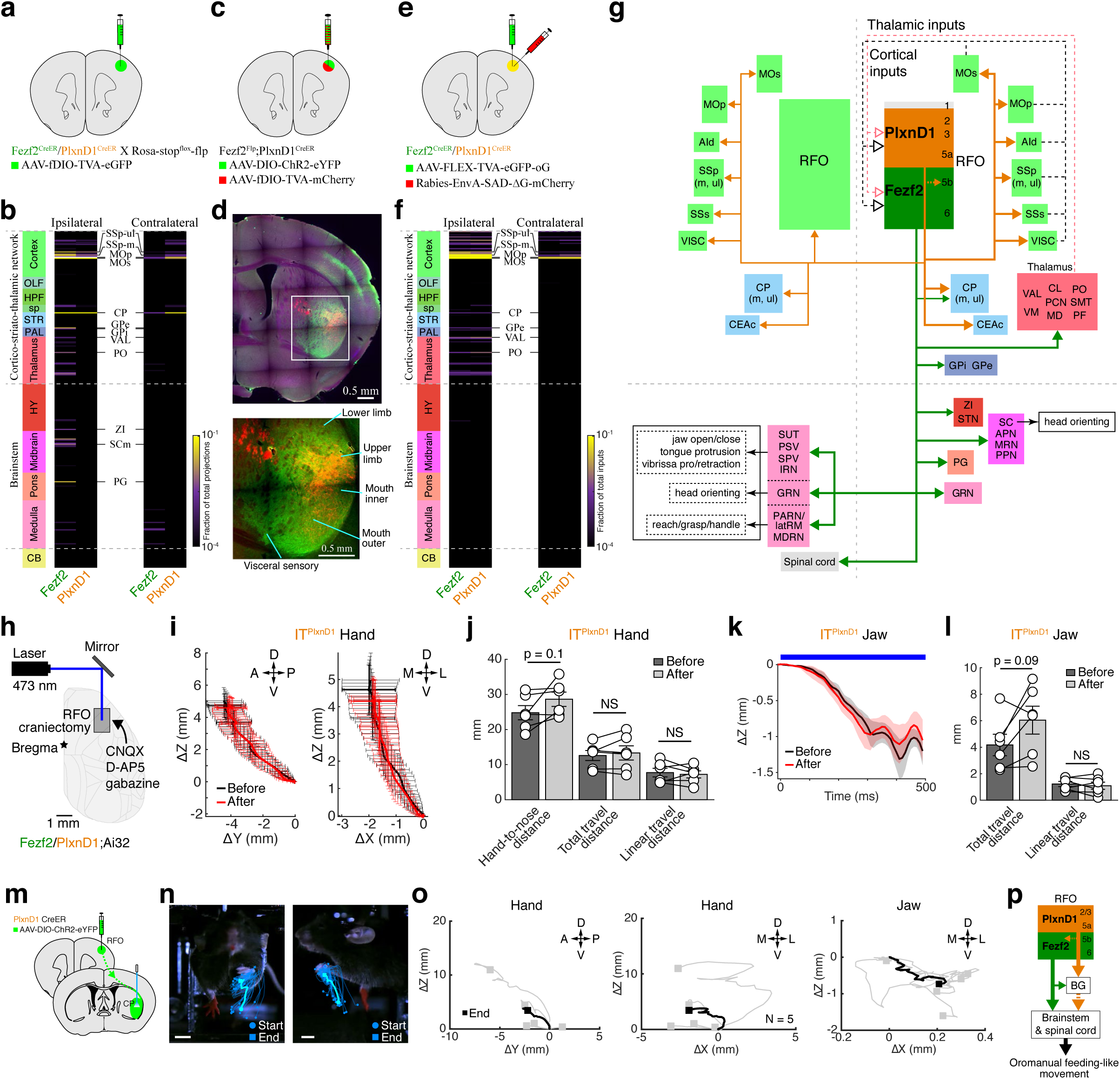
Input-output tracing of PTs^Fezf2^ and ITs^PlxnD1^ in RFO reveal a brain network for oromanual coordination. **a.** Schematic for anterograde tracing of PTs^Fezf2^ and ITs^PlxnD1^ in the RFO. **b.** Axon projection matrix from RFO to 315 ipsilateral and 315 contralateral targets (in rows), each grouped under 12 major categories (left column). Color shades in each column represent fraction of total axon signal averaged from 2 *Fezf2* and 2 *PlxnD1* mice. Related results are shown in **Extended Data Fig. 7**. **c.** Schematic for simultaneous anterograde tracing of PTs^Fezf2^ and ITs^PlxnD1^ in the RFO. **d.** RFO PTs^Fezf2^ show restricted projections (mCherry) whereas ITs^PlxnD1^ show broad projections (eYFP) in the striatum. Putative functional domains of the striatum were annotated (see Methods). **e.** Schematic for retrograde monosynaptic rabies tracing of PTs^Fezf2^ and ITs^PlxnD1^ in the RFO. **f.** Monosynaptic input matrix to RFO from 315 ipsilateral and 315 contralateral targets (in rows), each grouped under 12 major categories (left column). Color shades in each column represent fraction of total input cells averaged from 4 *Fezf2* and 5 *PlxnD1* mice. Related results are shown in **Extended Data Fig. 8**. **g.** A summary wiring diagram of efferent from (solid line) and afferent to (dashed line) PTs^Fezf2^ and ITs^PlxnD1^ in the RFO. Contralateral inputs were omitted for simplicity. Note that the IT^PlxnD1^ projection targets together delineate a bilateral RFO-centered cortico-striatal network, while PTs^Fezf2^ project unilaterally to a large set of subcortical targets, including brainstem areas implicated in both forelimb and orofacial actions. See text for detailed description. **h.** Schematic of optogenetic activation experiment with intracortical synaptic transmission blockade. **i.** 2D projections of left-hand trajectories following stimulation of RFO ITs^PlxnD1^. Black and red curves represent trajectories before and after drug application, respectively. **j.** Hand-to-nose distance, total- and linear-travel distance of hand movement before and after drug application. **k.** Movement trajectory of the jaw following stimulation of RFO ITs^PlxnD1^. Black and red curves represent trajectories before and after drug application, respectively. Blue bar represents stimulation window. Shading around mean denotes ± s.e.m. **l.** Total- and linear-travel distance of jaw movement before and after drug application. n = 6 mice in **i**-**l**. **m.** Schematic of AAV injection in the RFO of *PlxnD1* driver mice and optogenetic stimulation of IT^PlxnD1^ striatal axons. **n.** Example movement trajectories of the left hand (contralateral to the stimulation side). Light blue trajectories represent averages. Circle and square indicate start and end positions respectively. Scale bar, 5 mm. **o.** 2D projections of left-hand and jaw trajectories after stimulation of 5 *PlxnD1* mice. Black trajectories represent averages. Square indicates end position. Movement trajectories were normalized to the start position in **i**, **k**, **o**. Movement directions are indicated as: A, anterior; P, posterior; D, dorsal; V, ventral; M, medial; L, lateral. Data are mean ± s.e.m in **i**, **j**, and **l.** NS, not significant, two-sided paired t-test. **p.** A simplified schematic showing two key RFO output pathways that regulate oromanual movements, a PT^Fezf2^-mediated corticofugal pathway and an IT^PlxnD1^-mediated cortico-basal ganglia pathway. AId, agranular insular area, dorsal part; APN, anterior pretectal nucleus; BG, basal ganglia; CB, cerebellum; CEAc, central amygdalar nucleus, capsular part; CL, central lateral nucleus of the thalamus; CP, caudoputamen; GPe, globus pallidus, external segment; GPi, globus pallidus, internal segment; GRN, gigantocellular reticular nucleus; HPF, hippocampal formation; HY, hypothalamus; IRN, intermediate reticular nucleus; latRM, lateral rostral medulla; m, mouth; MD, mediodorsal nucleus of the thalamus; MDRN, medullary reticular nucleus; MOp, primary motor area; MOs, secondary motor area; MRN, midbrain reticular nucleus; OLF, olfactory areas; PAL, pallidum; PARN, parvicellular reticular nucleus; PCN, paracentral nucleus; PF, parafascicular nucleus; PG, pontine gray; PO, posterior complex of the thalamus; PPN, pedunculopontine nucleus; PSV, principal sensory nucleus of the trigeminal; SC, superior colliculus; SCm, superior colliculus, motor related; SMT, submedial nucleus of the thalamus; sp, cortical subplate; SPV, spinal nucleus of the trigeminal; SSp-m, primary somatosensory area, mouth; SSp-ul, primary somatosensory area, upper limb; SSs, secondary somatosensory area; STN, subthalamic nucleus; STR, striatum; ul, upper limb; SUT, supratrigeminal nucleus; VAL, ventral anterior-lateral complex of the thalamus; VISC, visceral area; VM, ventral medial nucleus of the thalamus; ZI, zona incerta.

In contrast, ITs^PlxnD1^ project bilaterally to primary and secondary motor (MOp, MOs) and sensory (SSp, SSs) orofacial (especially mouth) and forelimb (especially upper limb) areas, and to dorsal agranular insular cortex (AId), visceral cortex (VISC), and the capsular part of the central amygdala nucleus (CEAc) (**Fig. 3b, Extended Data Fig. 7b**, d**, Supplementary Video 6**). Furthermore, ITs^PlxnD1^ prominently project to bilateral ventrolateral striatum (**Fig. 3b, Extended Data Fig. 7b**, d), a region implicated in food handling and feeding ^53–55^. Simultaneous anterograde mapping of PT^Fezf2^ and IT^PlxnD1^ projections in the striatum of the same mouse revealed that whereas the former are highly focused to the upper limb and mouth domains, the latter are much more broad and additionally cover the lower limb and visceral sensory domains (**Fig. 3c, d**, **Extended Data Fig. 7e**). Together, these multi-region projection patterns suggest that RFO ITs^PlxnD1^ regulate a large set of corresponding cortical and striatal targets related to forelimb and oral sensorimotor as well as visceral and appetitive functions.

Retrograde monosynaptic rabies tracing revealed that cortical inputs to ITs^PlxnD1^ and PTs^Fezf2^ of the RFO are derived almost exclusively from their projection targets (i.e., forelimb and orofacial sensorimotor areas, AId, and VISC; **Fig. 3e, f, Extended Data Fig. 8a**, b, d, f**, Supplementary Video 6**). In addition, ITs^PlxnD1^ and PTs^Fezf2^ receive major subcortical inputs from the thalamus, including the ventral anterior-lateral complex and posterior complex (**Fig. 3f, Extended Data Fig. 8b**, e, f). Another robust subcortical input source is the external segment of the globus pallidus (**Extended Data Fig. 8c**). Together, these results demonstrate that RFO forms a reciprocally connected and thalamic driven cortical network involving the forelimb and orofacial sensorimotor areas as well as insular and visceral areas (**Fig. 3g**). Within the RFO, ITs^PlxnD1^ project bilaterally to all areas of this cortical network and their corresponding striatal targets, while PTs^Fezf2^ project unilaterally to a large set of subcortical targets implicated in both forelimb and orofacial control (**Fig. 3g**). Altogether, these findings delineate the organizational scheme and PN type implementation of an RFO-centered brain circuit poised to regulate concerted feeding actions involving oromanual coordination.

To explore the functional relationship between RFO PT^Fezf2^ and IT^PlxnD1^ output pathways in regulating forelimb and orofacial movements, we examined the effects of PT^Fezf2^ and IT^PlxnD1^ activation in head-fixed mice following blockade of intracortical synaptic transmission (**Fig. 3h**, **Extended Data Fig. 9a**). The hand and mouth movement trajectories induced by PT^Fezf2^ and IT^PlxnD1^ activation were largely unaffected by the application of glutamate and GABA_A_ receptor antagonist (CNQX, D-AP5, and gabazine) to the cortical surface (**Fig. 3i-l**, **Extended Data Fig. 9b-g**). Although this result does not exclude potential intracortical connectivity between PTs^Fezf2^ and ITs^PlxnD1^, it suggests that IT^PlxnD1^ activation can induce concurrent mouth and forelimb movements independent of local PT^Fezf2^ activation, likely through recruitment of the striatal targets of RFO ITs^PlxnD1^. Consistent with this notion, stimulating axon terminals of RFO ITs^PlxnD1^ in the ipsilateral striatum induced concurrent contralateral but not ipsilateral hand-to-mouth withdraw and mouth opening (**Fig. 3m-o**, **Extended Data Fig. 9h**). These results suggest that in addition to a local ITs^PlxnD1^ to PTs^Fezf2^ cortical connection ^33,56^ and PT^Fezf2^-mediated corticofugal pathway, ITs^PlxnD1^ can influence oromanual movement through the basal ganglia circuit (**Fig. 3p**).

### RFO PT^Fezf2^ and IT^PlxnD1^ activity correlate with pasta retrieval and oromanual manipulation

To examine whether and how PN type activity in the RFO correlate with different action motifs of pasta eating in free-moving mice, we used fiber photometry to record population calcium dynamics from PTs^Fezf2^ or ITs^PlxnD1^ in the right RFO, and, as a comparison, the left aCFA - an area involved in forelimb movement (**Fig. 4a, Extended Data Fig. 10a**). Interestingly, PT^Fezf2^ and IT^PlxnD1^ population activity patterns were broadly similar, we thus refer to them together as PN^Fezf2/PlxnD1^ activity. As mice entered the dining area (demarked by stepping across an elevated bar, **Fig. 2a**) to approach the pasta, PN^Fezf2/PlxnD1^ activity in aCFA was higher than that in RFO (**Fig. 4b-d**, **Extended Data Fig. 10b-d**). As mice licked to retrieve the pasta and then took a sitting posture and transferred the pasta from the mouth to the hands, RFO PN^Fezf2/PlxnD1^ activity sharply increased and then fell and rose in proportion to food handling vigor (**Fig. 4b, e, f, Extended Data Fig. 10b**, e, f). During the same period, aCFA PN^Fezf2/PlxnD1^ activity decreased to baseline levels (i.e., levels when mice were resting in the waiting area). After the pasta was consumed and as the mouse left the dining area, RFO activity dropped whereas aCFA activity increased (**Fig. 4b, Extended Data Fig. 10b**).

**Figure 4.**
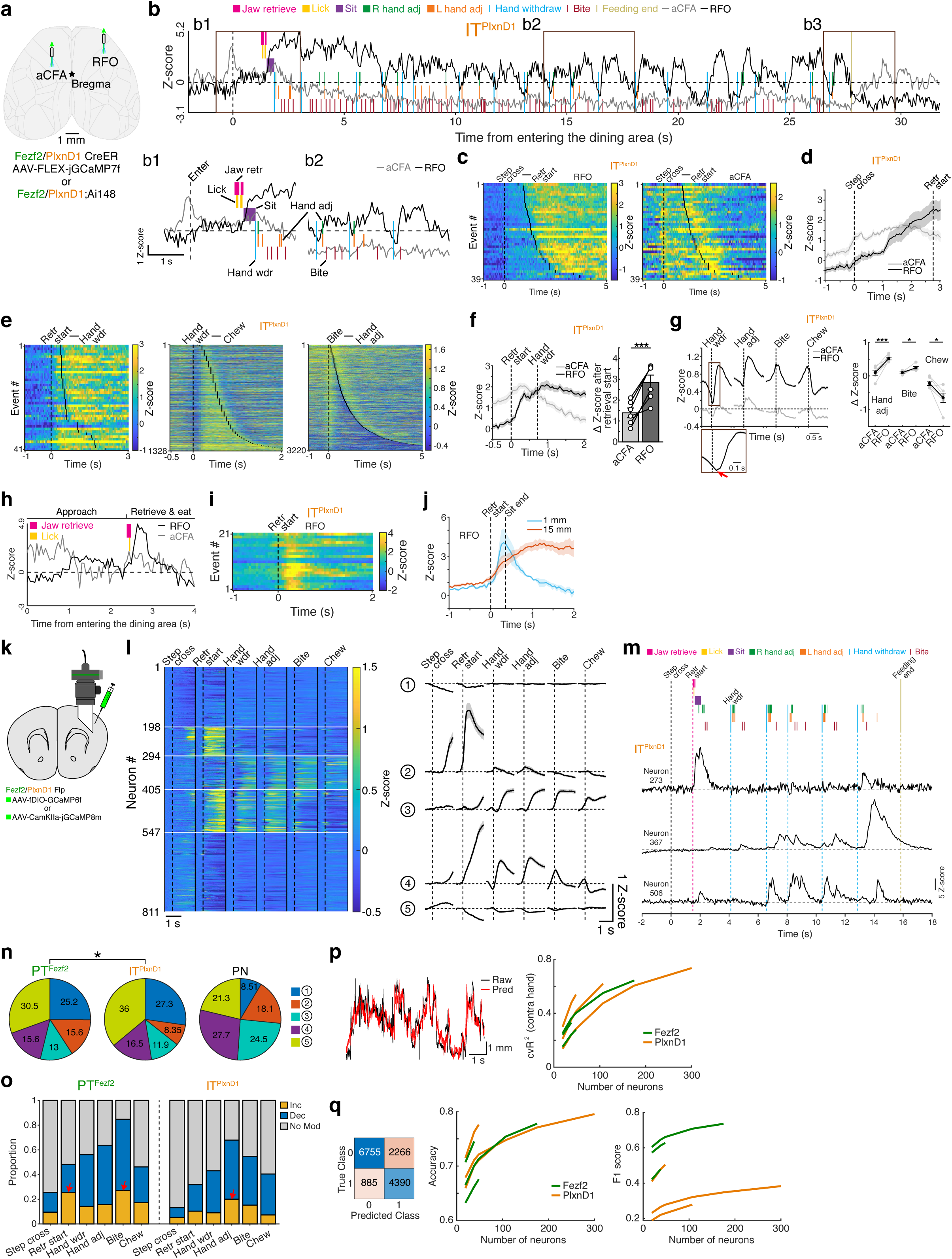
PT^Fezf2^ and IT^PlxnD1^ activity in RFO correlate with pasta retrieval and oromanual manipulation. **a.** Schematic depicting fiber photometry from the right RFO and left aCFA. **b.** Single-trial calcium activity traces of ITs^PlxnD1^ in the RFO (black) and aCFA (gray) of a mouse eating 15-mm angel-hair pasta overlaid on the actogram. Example time windows are highlighted (**b1-b3**) and two of them are expanded (**b1** and **b2)**. The rise of aCFA activity at time 0 (dashed line) correlates with crossing a step when entering the dining area (Fig. 2a). **c.** Heat maps of IT^PlxnD1^ population activity in the RFO and aCFA aligned to entry into the dining area and sorted by pasta retrieval start. **d.** Averaged population activity of ITs^PlxnD1^ in the RFO and aCFA aligned to entry into the dining area (n = 6 mice). Vertical dashed lines indicate average time to the retrieval start. **e.** Heat maps of RFO IT^PlxnD1^ population activity aligned to retrieval start (left), hand withdraw (middle), and bite (right). Activity traces were sorted by the earliest hand withdraw (left), chew (middle), and hand adjustment (right) events, respectively. **f, g.** Averaged IT^PlxnD1^ population activity in the RFO and aCFA aligned to retrieval start (**f**; left panel) and hand withdraw, hand adjustment, bite, and chew (**g**; left panel). Vertical dashed lines indicate average time to the first hand withdraw in **f**. Changes in population activity are shown in the right panels. RFO IT^PlxnD1^ activity rise after the onset of hand withdraw with a lag (red arrow in the expanded window of **g)**. n = 6 mice; *p < 0.05, ***p < 0.005, two-sided paired t-test. **h, i.** Single-trial IT^PlxnD1^ activity in the RFO and aCFA as a mouse consumed 1-mm angel-hair pasta without sitting up and hand recruitment (**h**). Key actions (colored annotations) were overlaid on the activity traces. Corresponding calcium activity (**i**) were aligned to retrieval start. **j.** Averaged RFO IT^PlxnD1^ population activity aligned to retrieval start for 15-mm and 1-mm angel-hair pasta (n = 3 mice). Vertical dashed lines indicate the average time to establish a sitting posture when eating 15-mm pasta. Activity remained high when mice handled and ate 15-mm pasta but declined when eating 1-mm pasta without sitting and hand recruitment. **k.** Schematic of viral injection and prism lens placement in the RFO for miniscope recording. **l.** Left, heat map of average neuronal activity aligned to different action motifs. Horizontal white lines in the left panel divide the five clusters. Vertical dashed lines indicate the onset of each action. Vertical solid black lines in the left panel separate the activity to different action motifs. Right, average activity of neurons in each of the five clusters aligned to different action motifs. Neurons were grouped using spectral clustering (see Methods). Dashed horizontal lines in the right panel depict Z-score of 0. **m.** Activity of three simultaneously recorded ITs^PlxnD1^ in an exemplar trial. The actogram is shown at the top. Horizontal dashed lines depict Z-score of 0. **n.** Proportion of PTs^Fezf2^, ITs^PlxnD1^, and PNs in the RFO assigned to the five clusters. *p < 0.05, Chi-square test. **o.** Proportion of PTs^Fezf2^ and ITs^PlxnD1^ that were modulated by different action motifs. The two red arrows of the left panel highlight the action motifs (retrieval start and bite) with larger proportion of PTs^Fezf2^ showing activity increase. The red arrow of the right panel highlights the action motif (hand adjustment) with the largest proportion of ITs^PlxnD1^ showing activity increase. **p.** Linear model decoding of Z-axis trajectory of the contralateral hand. Left panel shows example raw and predicted hand-movement trajectories. Right panel shows the cross-validated explained variance by including different numbers of neurons in the linear model. **q.** Linear model decoding of the occurrence of bite. Left panel shows a confusion matrix of true and predicted classes. Right panels show accuracy and F1 score by including different numbers of neurons in the linear model. N = 262 neurons from 3 mice for PTs^Fezf2^, N = 455 neurons from 3 mice for ITs^PlxnD1^, and N = 94 neurons from 2 mice for PNs in **l, n, o**. Data from each mouse were plotted in **p**, **q**. Shading around mean denotes ± s.e.m in **d, f, g, j,** and **l**. Data are mean ± s.e.m in **f, g**.

We next analyzed RFO PT^Fezf2^ and IT^PlxnD1^ activity patterns during the handle-bite phase and the chewing phase that were automatically identified by a hidden Markov model (**see Methods**). Elevated PN^Fezf2/PlxnD1^ activity was specifically correlated with handle-bite movements but not with removing the pasta from mouth or subsequent chewing (**Fig. 4e, g, Extended Data Fig. 10e**, g). In particular, PN^Fezf2/PlxnD1^ activity increase was best correlated with hand adjustments that advanced the pasta for a bite and hand-assisted pasta biting/snapping (**Fig. 4e, g, Extended Data Fig. 10e**, g). Notably, a cross-correlation analysis revealed that, at the start of each bout, the elevation of RFO activity reliably *followed* hand withdraw, measured as decreased hand-to-nose distance (**Extended Data Fig. 10h**, i), suggesting that PN^Fezf2/PlxnD1^ activity was not associated with initiating the hand withdraw. In addition, there was an increase in IT^PlxnD1^ activity followed hand withdraw with a shorter delay compared to that of PT^Fezf2^ (**Fig. 4g, Extended Data Fig. 10g**, j), indicating that IT^PlxnD1^ activity leads that of PT^Fezf2^ in regulating oromanual manipulation.

To further clarify whether RFO PN^Fezf2/PlxnD1^ activity is associated with oromanual coordination or with eating using the mouth, we fed mice short (1-mm long) pieces of pasta, which were eaten without sitting up and handling (**Fig. 4h**, **Extended Data Fig. 10k**). PN^Fezf2/PlxnD1^ activity rose immediately as the mice picked up the pasta by mouth but then quickly decreased to baseline with chewing (**Fig. 4h-j**, **Extended Data Fig. 10k-m**). Together, these results indicate that in the handle-eat stage, population activity is associated with coordinated hand and oral movements for positioning the pasta in the mouth and for hand-assisted biting with incisors ^36^. The population activity of RFO PT^Fezf2^ and IT^PlxnD1^ is not associated with discrete forelimb or oral movements such as removing pasta from mouth and chewing with molars ^36^.

Although the population activity patterns are broadly similar between PTs^Fezf2^ and ITs^PlxnD1^ in the RFO, single-neuron activity in the motor cortex are often heterogeneous and encode different aspects of movement ^57–59^. We therefore used miniscope recording to examine how single PN activity correlate with different action motifs of pasta eating (**Fig. 4k, Extended Data Fig. 10n**). In addition to PTs^Fezf2^ and ITs^PlxnD1^, we included a set of broad PNs infected by an AAV-CaMKII-GCaMP8m vector. The average activity of individual PTs^Fezf2^ and ITs^PlxnD1^ were highly consistent with the population activity acquired using fiber photometry (**Extended Data Fig. 10o**). There was a larger increase in average activity of PTs^Fezf2^ compared with ITs^PlxnD1^ during the entire handle-eat stage (**Extended Data Fig. 10o**). An unsupervised clustering analysis of neuronal activity in relation to different action motifs revealed five functional clusters with high stability for the cluster assignment (**Extended Data Fig. 10p-r**). Four of these clusters included neurons that increased their activity at different stages of the behavior (**Fig. 4l, m, Extended Data Fig. 10s**). Notably, cluster Ⅱ neurons greatly increased their activity during tongue-jaw mediated pasta retrieval but not other actions and so we term these “oral-retrieval cells”. Cluster III neurons exhibited elevated activity at bite and chew and so we term these “oral-eat cells”. Cluster Ⅳ neurons selectively increased their activity during oromanual-mediated hand adjustment and bite; we term these “oromanual cells”. This result suggests that there are distinct functional subsets of RFO PNs differentially involved in oral and oromanual actions. Notably, cluster Ⅱ contained a larger proportion of PTs^Fezf2^ than ITs^PlxnD1^ (**Fig. 4n**), suggesting that PTs^Fezf2^ play a more prominent role in pasta retrieval. In addition, the proportions of neurons with significantly modulated activity to different action motifs differed between PTs^Fezf2^ and ITs^PlxnD1^ (**Fig. 4o**). Whereas a larger proportion of PTs^Fezf2^ increased their activity at pasta retrieval and bite compared to other actions, the largest proportion of ITs^PlxnD1^ modulated their activity during hand adjustments (**Fig. 4o**).

To further examine the relationship between RFO neuronal activity with hand and mouth movements, we performed linear decoding analyses using activity of different numbers of neurons recorded in the same behavioral session. We found that PT^Fezf2^ and IT^PlxnD1^ activity decoded hand movement trajectory and the occurrence of bites with high accuracy (**Fig. 4p, q**). The decoding performance increased as more neurons were included in the model. This result strongly suggests that PTs^Fezf2^ and ITs^PlxnD1^ in the RFO contribute to the dexterous hand and mouth movements and the coordination between the two for eating.

### RFO PN types differentially regulate online food manipulation

Although PT^Fezf2^ and IT^PlxnD1^ activity patterns are broadly similar and both include the five functional cell clusters (**Fig. 4n**), they may nevertheless differentially regulate different components of the pasta-eating behavior through their highly distinct downstream circuits (**Fig. 3g**). Indeed, our optogenetic activation of RFO PTs^Fezf2^ and ITs^PlxnD1^ using the same stimulation pattern led to significantly different movement output (**Fig. 1e, k**). Thus, to examine when and how these different neuronal populations regulate real-time oromanual behavior, we optogenetically suppressed the activity of PTs^Fezf2^ and ITs^PlxnD1^, as well as the broad PNs^Emx1^ as a comparison, at different stages of the pasta-eating sequence (**Fig. 5a, Extended Data Fig. 11a**). Bilateral inhibition of the three populations during the retrieve-eat stage did not perturb approach and pasta detection but increased the overall feeding duration (**Fig. 5b, Extended Data Fig. 11b-d**). In particular, PN^Emx1^, PT^Fezf2^, and IT^PlxnD1^ inhibition all increased retrieval attempts (**Fig. 5b, c, Extended Data Fig. 11b**, e**, Supplementary Video 7**), likely due to impairments in tongue movement. Inhibition of PNs^Emx1^ or PTs^Fezf2^, but not ITs^PlxnD1^, delayed successful pasta retrieval (**Fig. 5b, d, Extended Data Fig. 11b**, f). Following pasta retrieval and transfer to the hands, inhibition of PNs^Emx1^ and ITs^PlxnD1^, but not PTs^Fezf2^, significantly increased the time taken to make the first bite (**Extended Data Fig. 11b**, g); this was due to difficulty in properly positioning the pasta inside the mouth, which requires coordinated hand and mouth movements including unimanual and bimanual hand adjustment (**Fig. 5e, Extended Data Fig. 11b**, h, i). These results suggest that whereas PTs^Fezf2^ play a more prominent role in dexterous tongue retrieval, ITs^PlxnD1^ play a more important role in oromanual coordination.

**Figure 5.**
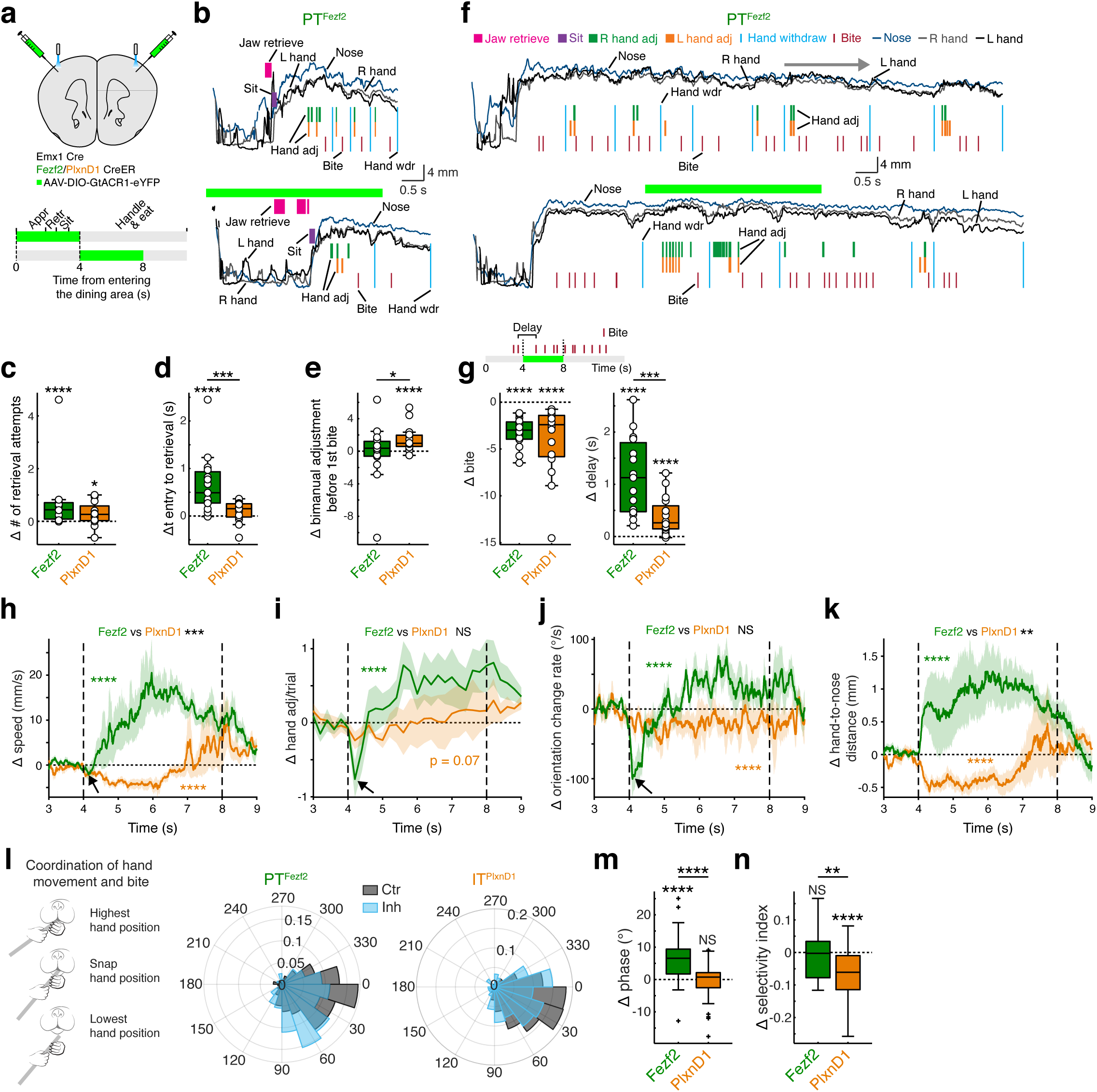
RFO PTs^Fezf2^ and ITs^PlxnD1^ regulate distinct components of oromanual manipulation. **a.** Schematic for optogenetic inhibition of PN types. AAV-DIO-GtACR1-eYFP was injected bilaterally into the RFO. Two inhibition schemes were directed to the retrieve-eat and handle-eat stages, respectively. Green bars indicate 4-s inhibition. **b.** Actograms of a mouse in control (upper) and PT^Fezf2^ inhibition (lower; green bar) trials. Z-axis trajectories of nose, right and left hands are shown. Actions are colored and labeled. **c, d.** PT^Fezf2^ inhibition interfered with pasta retrieval, measured as increased number of retrieval attempts (i.e., retrieval jaw movements) (**c**) and lengthened the time from entry to successful retrieval (**d**). **e.** IT^PlxnD1^ inhibition increased bimanual adjustment before the first bite. **f.** Actograms of a mouse at the handle-eat stage in control (top) and PT^Fezf2^ inhibition (bottom) trials. Z-axis trajectories of nose and two hands are shown. The top horizontal arrow indicates a control handle-eat bout. PT^Fezf2^ inhibition led to substantially increased hand adjustments. **g.** PT^Fezf2^ and IT^PlxnD1^ inhibition resulted in decreased number (left) and increased delay (right) of bites. Red ticks in the top schematic indicate bite events. **h-k**. Temporal changes of Z-axis hand speed (**h**), number of hand adjustment (**i**), pasta-orientation change rate (**j**), and hand-to-nose distance (**k**) during PT^Fezf2^ and IT^PlxnD1^ inhibition in the handle-eat stage. The angle between the pasta and the XY plane was measured to compute the pasta-orientation change rate (see **Extended Data Fig. 11m**). Note that hand speed, number of hand adjustment, and pasta-orientation change rate decreased briefly at the beginning of the PT^Fezf2^ inhibition (black arrows in **h-j**) followed by an increase. Vertical dashed lines indicate the 4-8 s inhibition. Shading around mean denotes ± s.e.m. n = 7 mice for PTs^Fezf2^ or ITs^PlxnD1^. **p < 0.01, ***p < 0.005, ****p < 0.001, two-way repeated measures ANOVA for the comparison of Control vs Inhibition and PT^Fezf2^ vs IT^PlxnD1^ data. **l.** Probability distributions of the phases of support-hand movement at the time of bites in control and PT^Fezf2^ or IT^PlxnD1^ inhibition trials. The narrower the probability distribution the higher the selectivity index. Schematic in the left panel depicts the coordination of downward hand movement with bite to snap the pasta. **m**, **n**. Average hand-movement phase at the time of a bite (**m**) and selectivity index of the bite-related hand-movement phases (**n**). Results from support and guide hands were similar and thus were pooled. n = 14 mice for PTs^Fezf2^ and 15 mice for ITs^PlxnD1^. Analyses in **g-n** were carried out for the same 4-8 s window for control and inhibition trials. n = 15 mice for PTs^Fezf2^ and 16 mice for ITs^PlxnD1^, for the analyses in **c-e, g**. *p < 0.05, **p < 0.01, ***p < 0.005, ****p < 0.001, two-sided paired t-test and two-sided Wilcoxon signed-rank test for the comparison of Control vs Inhibition, two sided t-test and two-sided Wilcoxon rank sum test for the comparison of effect size between PT^Fezf2^ and IT^PlxnD1^ inhibition in **c-e, g, m, n**. NS, not significant.

When we optogenetically suppressed these neuronal populations during the handle-eat stage, we observed significant increases in eating duration, delays in pasta biting, and reductions in overall pasta consumption (**Fig. 5f, g, Extended Data Fig. 11j-l**). The bite delay was significantly longer during PT^Fezf2^ than IT^PlxnD1^ inhibition, consistent with an initial pause of hand and mouth movements during PT^Fezf2^ inhibition (**Fig. 5g-i**). These deficits were not due to an impairment in biting *per se*. When we presented pasta in a holding device where mice could bite without using their hands, PN inhibition did not interfere with pasta biting (**Extended Data Fig. 12, Supplementary Video 8)**. Notably, broad PN^Emx1^ inhibition in four of the six mice prevented their proper handling and positioning of the pasta for a single bite (**Extended Data Fig. 11j**, l). The initial pause of oromanual movements caused by PT^Fezf2^ inhibition was followed by many uncoordinated hand movements including hand adjustment, leading to frequent pasta-orientation changes that were ineffective for pasta biting (**Fig. 5h-j, Supplementary Video 9**). These uncoordinated hand movements also resulted in an increased distance between the hand and mouth (**Fig. 5k**). In contrast, IT^PlxnD1^ inhibition led to stiffness of forelimb movement and reduced pasta manipulation, manifested as a prolonged decrease in hand movement speed, pasta orientation changes, and hand-mouth distance (**Fig. 5h, j, k, Supplementary Video 9**). These findings suggest that whereas PTs^Fezf2^ play a more prominent role in pasta positioning, ITs^PlxnD1^ are more involved in coordinating the oromanual movements required for a bite. These difficulties in pasta manipulation were further confirmed by more variable (PNs^Emx1^) and more vertical orientations (PTs^Fezf2^ and ITs^PlxnD1^) of pasta that deviated from the bite-effective orientation during the inhibition (**Extended Data Fig. 11m**, n).

These manipulations revealed differential impairments in the mechanics of pasta manipulation during the inhibition of specific neuronal subtypes, even when mice occasionally managed to bite/snap the pasta. While mice normally snap the pasta within a characteristic phase range of the downward hand-swing, PT^Fezf2^ inhibition resulted in a lower hand-swing position at the moment of a bite (i.e. increased phase), indicative of a less effective hand and/or jaw movements for snapping (**Fig. 5l, m**). IT^PlxnD1^ inhibition led to an increased variability of hand-swing position at the moment of a bite, suggesting a deficit in the precise temporal coordination between hand and jaw movements for pasta snapping (**Fig. 5l, n**). Altogether, our results indicate that PNs^Emx1^, PTs^Fezf2^, and ITs^PlxnD1^ in the RFO regulate the execution and coordination of oromanual manipulation in using the hands to position the pasta in the mouth and then exert force on it to snap the pasta in coordination with a bite. Whereas PTs^Fezf2^ are preferentially involved in regulating dexterous hand and mouth movements, ITs^PlxnD1^ play a more prominent role in coordinating oromanual manipulation during pasta biting/snapping.

## DISCUSSION

The sophisticated coordination of hand and mouth movements for food manipulation is distinctive in rodents and primates and features a number of action motifs that comprise the syllables of eating behavior ^5,43^. Through a comprehensive approach integrating high-speed videography, sound recording, and quantitative behavioral analysis in freely moving mice, we have characterized the sequential stages and multiple action motifs of pasta eating—with particular emphasis on the precise coordination between the hand and mouth movements for pasta manipulation and biting. Combining systematic optogenetic screen of cortical areas and projection neuron (PN) types, cell-type resolution input-output circuit tracing, neural recording, and functional manipulation in freely feeding mice, we discovered a cortical motor area and identified the underlying cell-type-specific circuit architecture that orchestrate oromanual food manipulation.

In relation to previously identified motor areas, RFO is localized to the posterolateral part of the recently delineated lateral anterior cortex (LAC) ^9^, a large territory containing subregions implicated in both skilled forelimb ^9^ and orofacial ^41,60^ movement. In particular, the tjM1 was identified in a whisk-to-lick task in which forelimb movement was not examined ^41^. We found that RFO has considerable overlap with tjM1. Thus, RFO is juxtaposed between a classically identified forelimb (CFA) ^21^ and an oral motor area (tjM1). This location likely optimizes the interaction between the oral and forelimb motor regions, with a plausible evolutionary implication that could be examined in developmental and comparative studies. It is noteworthy that stimulation of the macaque precentral gyrus, a region similarly juxtaposed between mouth and hand motor areas, induces concomitant oromanual movements ^15,61^. Similarly, the human precentral gyrus contains neurons that respond to mouth stimuli and elicit concurrent hand-to-mouth and mouth movements when stimulated ^62^. Together, these findings suggest that a conserved RFO homologue contributes to food manipulation behaviors in rodents and primates including humans.

The motor cortex orchestrates movement through several major pathways that engage other cortical areas and many subcortical structures ^63–65^. Our PN-type based antero- and retro-grade tracing delineates a scheme of cell type implementation of an RFO-centered circuit organization comprising several connectivity components. First, PTs^Fezf2^ project unilaterally to a large set of subcortical targets, including the thalamus, striatum, lateral superior colliculus, pons, medulla, and spinal cord. These targets include areas for both forelimb and orofacial actions such as reaching ^8,12^, grasping ^8,12,48^, jaw opening ^11,49,50^, and licking ^11,50,51^ (**Fig. 3g**). Consistent with these projections, PT^Fezf2^ activation induced contralateral and relatively limited head-forelimb-orofacial movements. Future single-cell reconstruction analysis ^66^ may reveal whether the innervation of both oral and forelimb targets is achieved by two separate PT^Fezf2^ subpopulations or by the collaterals of individual neurons. In contrast, ITs^PlxnD1^ project bilaterally to multiple cortical and striatal regions: the cortical targets include the motor (MOp, MOs) and sensory (SSp, SSs) areas of mouth and upper limb, while the striatal targets include the forelimb and oral domains of the ventrolateral striatum that has been implicated in food handling ^53–55^. Notably, these striatal domains targeted by RFO ITs^PlxnD1^ also receive topographically organized inputs from the cortical sensorimotor areas where RFO ITs^PlxnD1^ project to ^53^. Therefore, these RFO IT^PlxnD1^ projections delineate a coherent RFO-centered cortico-striatal network poised to regulate oral and manual coordination (**Fig. 3g**). Consistent with this scheme, IT^PlxnD1^ activation elicited bilateral and concerted head, body, orofacial, forelimb, and hand movements resembling eating; additionally, activation of the striatal branch of IT^PlxnD1^ axons induced concurrent mouth and forelimb movements. Considering that a major cortical PN connectivity motif is that ITs are generally upstream of PTs ^33^, these results suggest a circuit organization whereby RFO ITs^PlxnD1^ activate ventrolateral striatum to engage an oromanual feeding program as well as excite local PTs^Fezf2^ that directly broadcast output throughout relevant subcortical regions (**Fig. 3p**). In parallel, through their intracortical projections (**Fig. 3g**), ITs^PlxnD1^ may concurrently influence the output of oral and forelimb related sensorimotor areas, which together coordinate oral and manual actions. Notably, RFO ITs^PlxnD1^ further project to the insular and visceral cortices and the capsular part of the central amygdala nucleus, areas implicated in appetitive behavior ^67–69^, suggesting an even higher level of coordination between the sensorimotor and valence systems.

Superimposed upon this output organization, our synaptic input tracing revealed that RFO ITs^PlxnD1^ and PTs^Fezf2^ receive cortical inputs exclusively from their projection targets described above, thereby forming an RFO-centered reciprocal cortical network, as well as inputs from various thalamic nuclei (e.g. the ventral anterior-lateral complex and posterior complex). These cortical and thalamic inputs may convey feedback and feedforward multi-sensory and movement state information to the RFO to regulate the online coordination of oromanual actions. Taken together, these input-output connectivity patterns delineate an RFO cortical circuit and associated brain network that orchestrate multi-body-part actions and coordinate oromanual manipulation.

Our quantitative analyses of the distinct stages and action motifs of the pasta eating behavior set the stage for exploring the neural control mechanisms of oromanual manipulation. We consider that the most striking feature of RFO PT^Fezf2^ and IT^PlxnD1^ population activity is their tight correlation with cooperative oromanual actions rather than with discrete forelimb or oral actions. In contrast, activity in the aCFA, a forelimb motor area implicated in forelimb reaching and locomotion behaviors ^7,21,24,70,71^, increases during the approach-to-retrieve stage but decreases during the handle-eat stage, a largely complementary pattern to that of RFO. The lag between the rise of RFO PN^Fezf2/PlxnD1^ activity and the initiation of hand withdraw is consistent with the finding that activity in the medial forelimb M1 but not lateral M1 (a region that overlaps with RFO) is related to hand transport of food to the mouth ^7^, suggesting that RFO activity does not initiate the hand withdraw. Furthermore, our cellular-resolution recordings reveal functional subsets of PNs that are selectively active during oral retrieval (tongue lick and incisor grasp), oral eating (bite and chew), and oromanual (hand adjustment and pasta snap) actions. Notably, the proportion of oromanual cells was the highest among these three subsets. While our analysis uncovered relatively small differences in activity patterns between PTs^Fezf2^ and ITs^PlxnD1^, this is perhaps not surprising given their similar synaptic input source (**Fig. 3g**) and the limited temporal resolution of calcium indicators. Similar activity patterns in PTs^Fezf2^ and ITs^PlxnD1^ could nevertheless mediate different circuit functions through distinct projection targets, as demonstrated by the different movement outcomes induced by the same optogenetic stimulation parameters (**Fig. 1e, k**). Future large-scale ^72^ and cell-type resolution ^73^ electrophysiology recordings in the RFO during pasta eating may reveal additional neuronal activity patterns and more distinctions between PN types.

Optogenetic inhibition of PTs^Fezf2^ and ITs^PlxnD1^ during different stages of pasta eating reveal their distinct roles in oromanual manipulation and pasta eating. During the retrieve-eat stage, PT^Fezf2^ inhibition interfered with pasta retrieval, likely due to impairments in dexterous tongue movement. IT^PlxnD1^ inhibition increased hand adjustment and thus impaired the coordination between the hand and mouth in passing the pasta between them for a first bite. During the handle-eat stage, whereas PT^Fezf2^ inhibition resulted in an initial pause of oral and manual movements followed by excessive and uncoordinated hand adjustments, IT^PlxnD1^ inhibition led to stiffness of hand movement and reduced pasta manipulation. These results align with our recent study in head-fixed mice that PT^Fezf2^ inhibition in an area overlapping with RFO impaired pellet insertion into the mouth, while IT^PlxnD1^ inhibition resulted in rigid pellet holding using digits ^74^. In addition, our hand-movement phase analysis revealed that, at the time of pasta biting, IT^PlxnD1^ inhibition increased variability of hand-swing position, whereas PT^Fezf2^ inhibition resulted in a lower hand-swing position. The former impairment suggests a disruption of the precise timing between hand and mouth movements and the latter impairment indicates less effective hand and/or mouth movements in pasta snapping. These results indicate that whereas PTs^Fezf2^ are preferentially involved in regulating dexterous hand and mouth movements, ITs^PlxnD1^ play a more prominent role in coordinating oromanual movements during pasta biting/snapping. Notably, RFO PN inhibition did not completely abolish hand movements or oromanual manipulation, likely due to the fact that IT^PlxnD1^ is a subset of the whole IT class and that a distributed cortical network involving multiple other brain areas contributes to the oromanual feeding behavior. Considered together with the distinct output circuits (**Fig. 3g**) and activation-induced movements of PTs^Fezf2^ and ITs^PlxnD1^ (**Fig. 1e, k**), our findings suggest that ITs^PlxnD1^ may coordinate actions across the body, including oromanual manipulation, for feeding, whereas PTs^Fezf2^ may predominantly initiate and regulate the dexterous oral and forelimb movements.

In summary, we discovered a cortical area and its key constituent cell types and their associated neural circuitry that orchestrate oromanual food manipulation during natural feeding behavior. Beyond cortex-centered studies, future work could examine how brain-wide neural communication ^75,76^ control and coordinate oromanual manipulation using large-scale neuropixels recording ^72^. The neural circuit for oromanual food manipulation revealed here may contribute to many other coordinated hand and mouth movements in rodents (e.g., pup cleaning, nest building) and humans (e.g., the coordination of hand and mouth in gesture associated speech) ^1,77^.

## ACKNOWLEDGEMENTS

We thank Yongjun Qian and Shengli Zhao for making the GtACR1 AAV vector and viruses; Joshua Hatfield for helping with STP imaging and video annotations; Priscilla Wu, Joshua Hatfield, and Bao-Xia Han for animal preparation and maintenance; Robert Eifert at the machine shop of CSHL for the fabrication of custom-designed parts; Weixin Zhong for drawing schematics; and Shiyang Pan, William Galbavy, Jason Tucciarone, Shreyas M. Suryanarayana, Patrick Mulcahey, and John Pearson for helpful discussions. We thank Richard Mooney and Nuo Li for comments on the manuscript. This work was supported in part by NIH grant U19MH114823-01 to Z.J.H. Z.J.H is also supported by a NIH Director’s Pioneer Award 1DP1MH129954-01.

## AUTHOR CONTRIBUTIONS

X.A. and Z. J. H. conceived this study. Z.J.H. supervised the study. X.A. designed the research and performed the majority of the experiments, and analyzed data. Y.L. performed in vivo electrophysiology recording. K.M. performed STP imaging and anatomy analyses. H.M. provided advice for data analysis. X.H.X. analyzed behavioral videos. A.K. and I.Q.W. made contributions to data analysis and discussion. Z.J.H. and X.A. wrote the manuscript with inputs from I.Q.W. and A.K.

---

**Extended Data Fig. 1.**
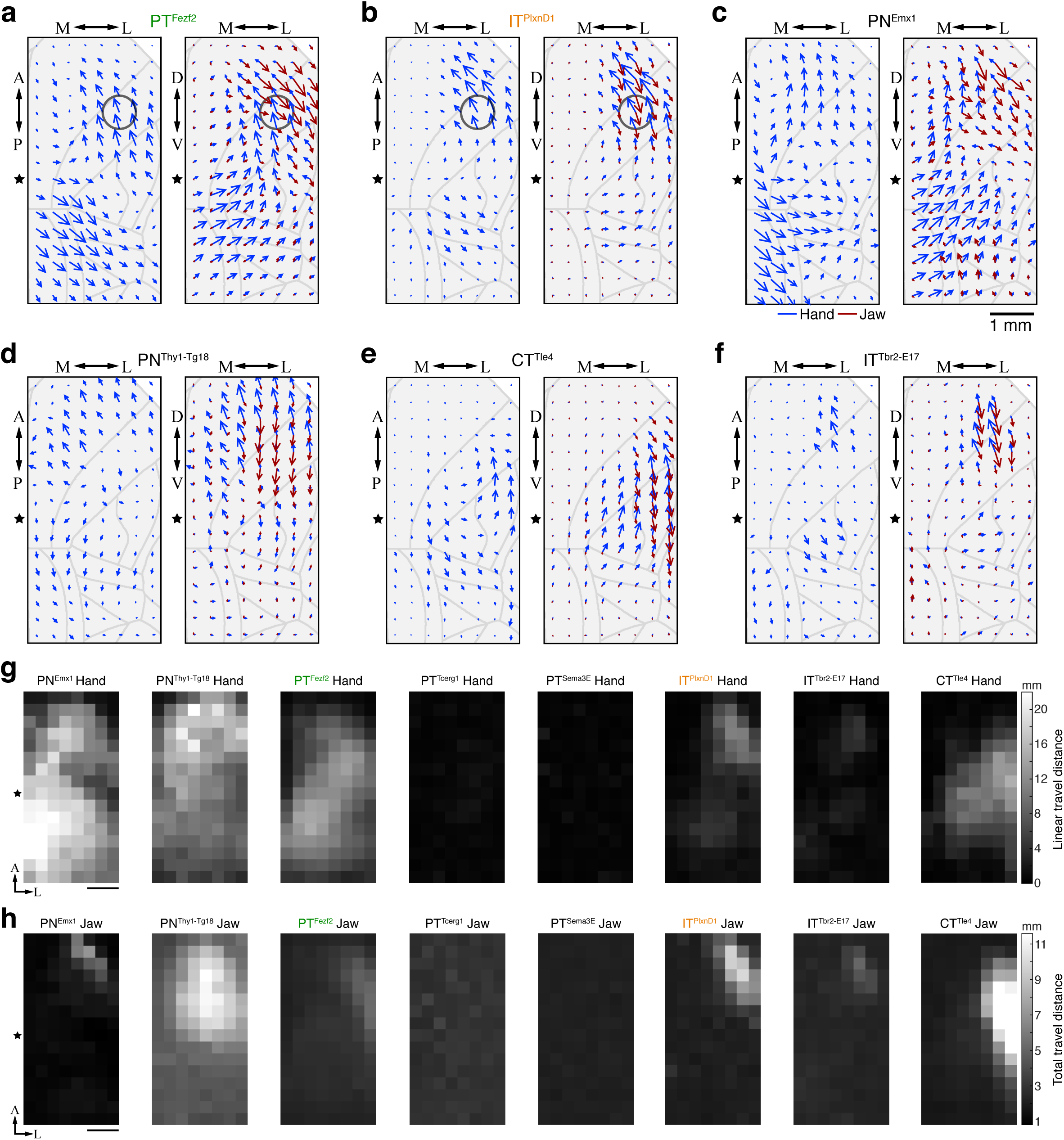
Different PN subpopulations exhibit distinct motor maps. Related to Fig. 1 **a-f.** Vector maps of hand (blue arrows) and jaw (red arrows) movement direction and distance following optogenetic activation of PTs^Fezf2^ (**a**), ITs^PlxnD1^ (**b**), PNs^Emx1^ (**c**), PNs^Thy1-Tg18^ (**d**), CTs^Tle4^ (**e**), and ITs^Tbr2-E17^ (**f**) in different locations of the dorsal cortex. Movement direction and distance along each axis are represented by arrow direction and length, respectively. Movement directions are indicated as: A, anterior; P, posterior; D, dorsal; V, ventral; M, medial; L, lateral. Distance was averaged across mice and normalized. n = 7 *PlxnD1*, 2 *Emx1*, 2 *Thy1-Tg18*, 5 *Tle4*, and 4 *Tbr2-E17* mice for the hand and jaw vector maps. n = 13 and 11 *Fezf2* mice for the hand and jaw vector maps respectively. **g.** Maps of hand movement distance (linear travel distance, measured from start to end). No clear hand movement was induced in *Tcerg1l* and *Sema3E* mice (Maps averaged from 2 *Emx1*, 2 *Thy1-Tg18*, 13 *Fezf2,* 5 *Tcerg1l*, 5 *Sema3E*, 7 *PlxnD1*, 4 *Tbr2-E17*, and 5 *Tle4* mice). **h.** Maps of total jaw movement distance. No clear jaw movement was induced in *Tcerg1l* and *Sema3E* mice (Maps averaged from 2 *Emx1*, 2 *Thy1-Tg18*, 11 *Fezf2*, 5 *Tcerg1l*, 5 *Sema3E*, 7 *PlxnD1*, 4 *Tbr2-E17*, and 5 *Tle4* mice). Stars indicate Bregma. Scale bars, 1 mm. Note, the cortical sites that induced hand and jaw movements by CT^Tle4^ activation were distinct and in a lateral region relative to the Bregma. This region aligns well to the primary sensory cortex of the upper limb (SSp-ul). CTs^Tle4^ in SSp-ul might possibly induce oromanual movements by activating motor cortical regions through a disynaptic cortico-thalamo-cortical pathway.

**Extended Data Fig. 2.**
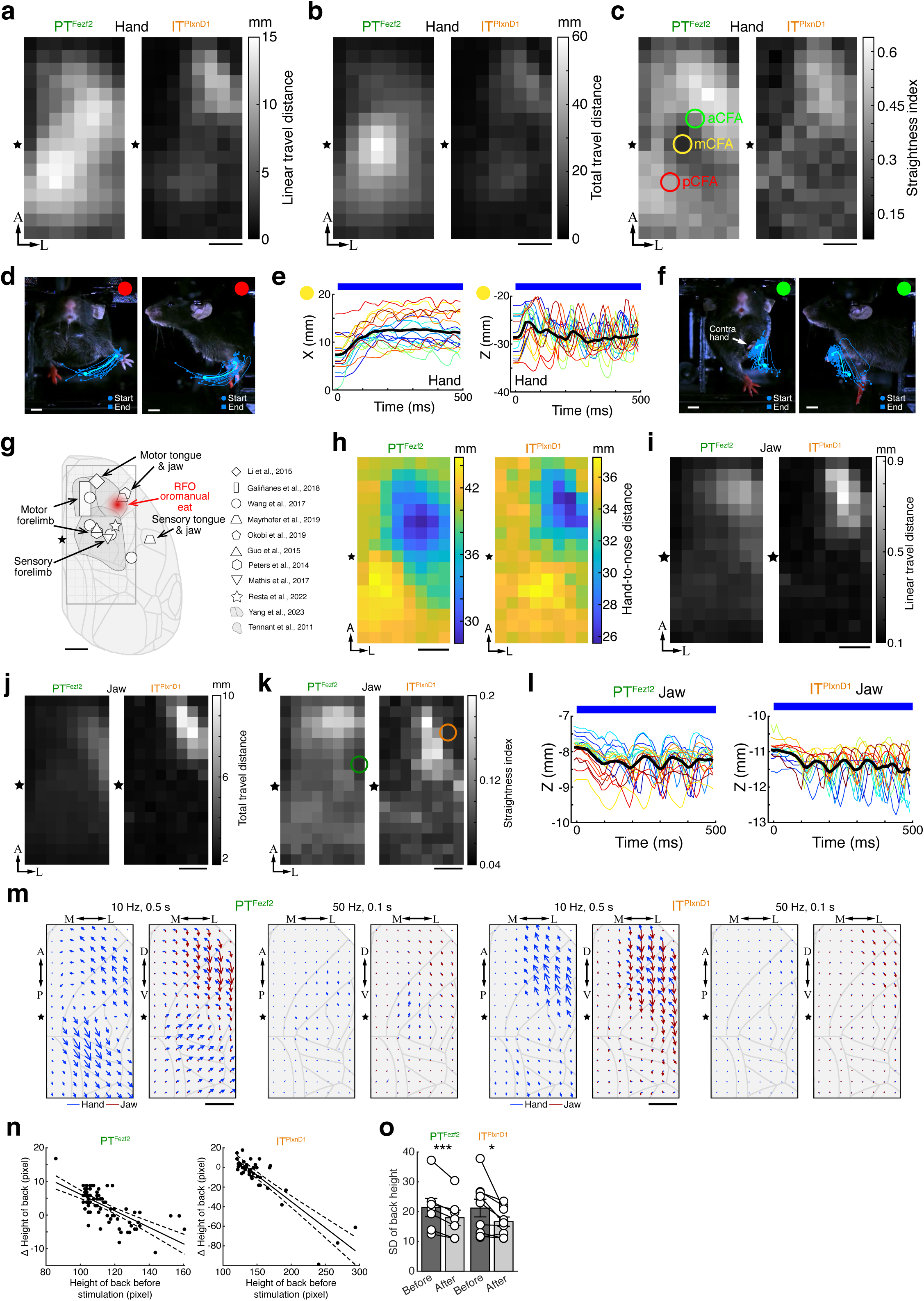
Characterization of forelimb and jaw movements induced by optogenetic activation of PTs^Fezf2^ and ITs^PlxnD1^. Related to Fig. 1. **a-c.** Maps of hand linear travel distance measured from start to end (**a**), total travel distance (**b**), and straightness index (**c**) (straightness index = linear travel distance/total travel distance, with smaller index = more rhythmic movement). **d-f.** Hand trajectories following PT^Fezf2^ activation at three sites as indicated by the three circles in **c.** Red circle at pCFA and 15 trials (**d**); yellow circle at mCFA and trajectory graphs of 19 trials showing repetitive movements (**e**); green circle at aCFA and17 trials (**f**). Light blue trajectories represent averages in **d**, **f**. Black trajectories in **e** indicate averages. The circle and square indicate start and end positions respectively in **d, f**. Note: the left hand is lifted and open after stimulation (white arrow in **f**). **g.** Schematic of RFO location in relation to other sensorimotor areas identified in previous studies (symbols and references). The grid indicates the area that was motor mapped in this study. **h.** Maps of hand-to-nose distance after activation. **i-k.** Maps of jaw linear travel distance measured from start to end (**i**), total travel distance (**j**), and straightness index (**k**) (Straightness index = linear travel distance/total travel distance, with smaller index = more rhythmic movement). **l.** Example jaw movement trajectories following PT^Fezf2^ or IT^PlxnD1^ activation at two sites as indicated by the two circles in **k** (20 PT^Fezf2^ and 16 IT^PlxnD1^ trials respectively). Black trajectories indicate averages. **m.** Vector maps of hand and jaw movement direction and distance acquired by optogenetic activation of PTs^Fezf2^ and ITs^PlxnD1^ using different stimulation parameters (compare with maps of 50 Hz, 0.5 s stimulation in **Extended Data Fig. 1b**, c). Movement direction and distance along each axis are represented by arrow direction and length, respectively. Movement directions are indicated as: A, anterior; P, posterior; D, dorsal; V, ventral; M, medial; L, lateral. Distance was averaged across mice and normalized to that from 10 Hz, 0.5 s stimulation (10 Hz, 0.5 s: n = 5 mice for PTs^Fezf2^ or ITs^PlxnD1^; 50 Hz, 0.1 s: n = 4 mice for PTs^Fezf2^ or ITs^PlxnD1^). **n.** Relationship between body height, measured as height of back of mice, before stimulation and change of body height after stimulation. Black solid line indicates linear fit (slope: -0.218 ± 0.061, R-squared: 0.36 ± 0.13 for PT^Fezf2^; slope: -0.307 ± 0.157, R-squared: 0.51 ± 0.17 for IT^PlxnD1^). Dashed lines depict 95% confidence bounds. **o.** Standard deviation of body height before and after stimulation. Data from 7 hemispheres of 5 mice and 9 hemispheres of 7 mice for PTs^Fezf2^ and ITs^PlxnD1^ respectively. Data are mean ± s.e.m. *p < 0.05, ***p < 0.005, two-sided paired t-test. Maps were averaged for 13 *Fezf2* and 7 *PlxnD1* mice in **a-c, h**, and 11 *Fezf2* and 7 *PlxnD1* mice in **i-k**. Blue bars in **e**, **l** represent stimulation window. Stars indicate Bregma. Scale bars, 1 mm in **a-c, g-k, m**; 5 mm in **d, f**.

**Extended Data Fig. 3.**
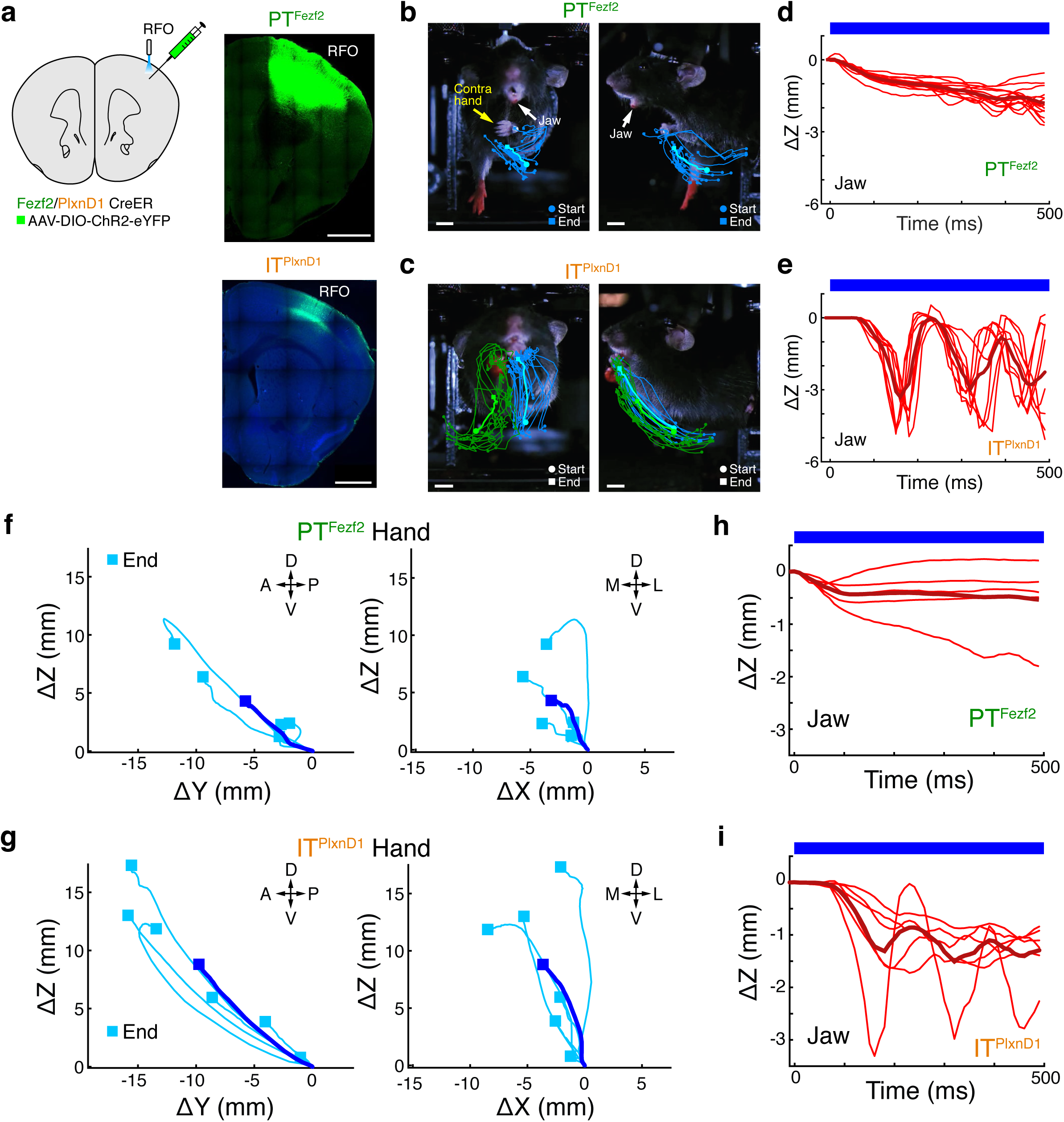
Activating AAV-targeted PTs^Fezf2^ or ITs^PlxnD1^ in RFO induces hand-to-mouth and mouth movements. Related to Fig. 1. **a.** Schematic of the approach and images of coronal sections showing PTs^Fezf2^ and ITs^PlxnD1^ infected by AAV-DIO-ChR2-eYFP injected into the right RFO. Scale bar, 1 mm. **b, c.** Example movement trajectories of the left hand (20 PT^Fezf2^ trials, **b**) and both hands (19 IT^PlxnD1^ trials, **c**). Light blue and green trajectories represent averages. Circles and squares indicate start and end positions respectively. Note: left hand is closed and jaw opens to the contralateral side after stimulation in **b.** Scale bar, 5 mm. **d, e.** Movement trajectories of the jaw (16 PT^Fezf2^ trials, **d**; 8 IT^PlxnD1^ trials, **e**). Note that whereas PT^Fezf2^ stimulation induced mostly jaw open, IT^PlxnD1^ stimulation induced rhythmic jaw movement. **f, g.** 2D projections of left-hand trajectories following stimulation in 5 *Fezf2* mice (**f**) and 6 *PlxnD1* mice (**g**). Square indicates end position. A, anterior; P, posterior; D, dorsal; V, ventral; M, medial; L, lateral. **h, i.** Movement trajectories of the jaw following stimulation in 5 *Fezf2* mice (**h**) and 6 *PlxnD1* mice (**i**). Movement trajectories were normalized to the start position in **d-i**. Dark red and blue trajectories represent averages in **d-i**. Blue bar in **d, e**, **h, i** represents stimulation window.

**Extended Data Fig. 4.**
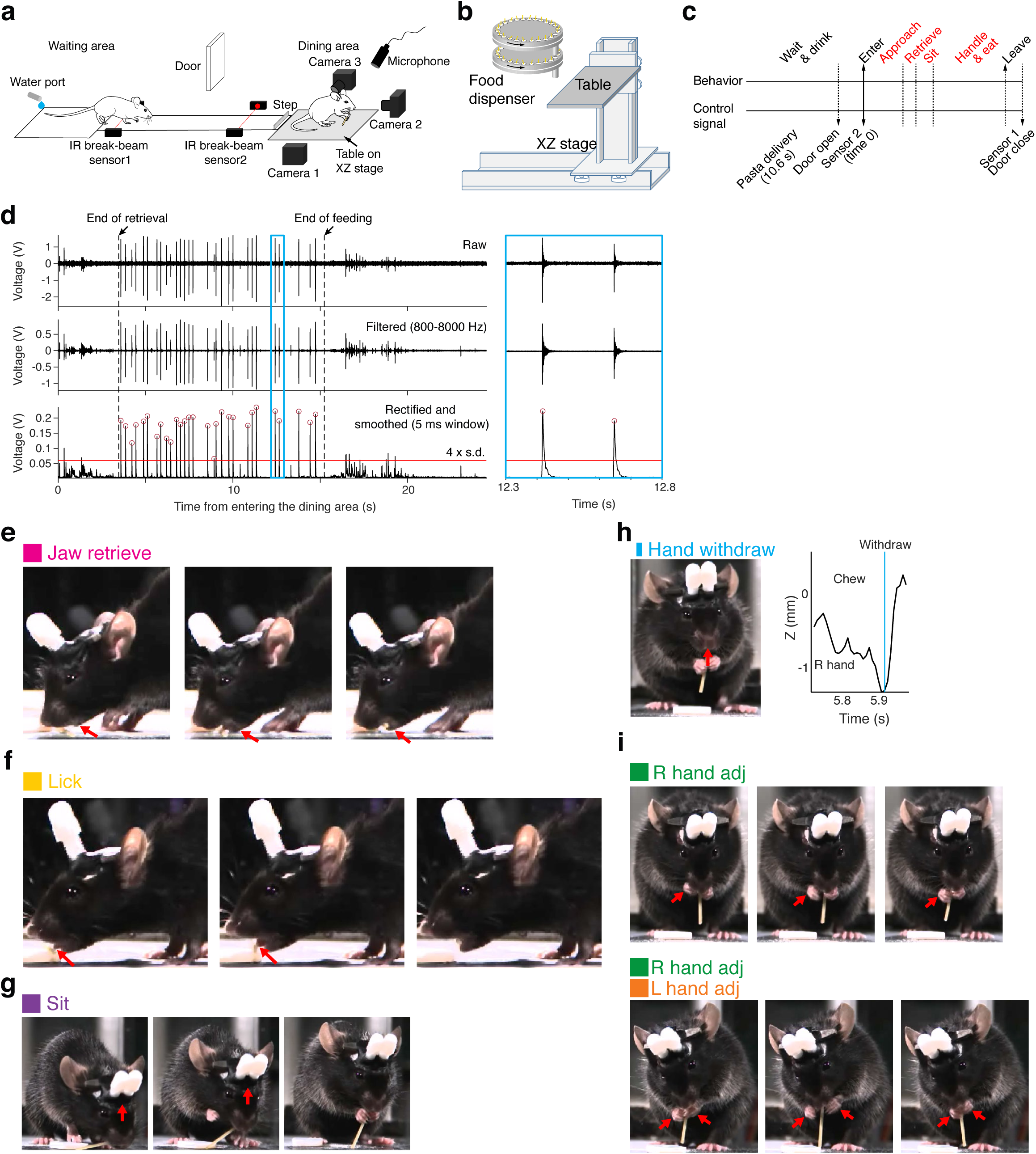
Design of the Mouse Restaurant and identification of action motifs in pasta eating. Related to Fig. 2. **a, b.** Schematic of the Mouse Restaurant. A table mounted on an XZ stage (**b**) brings food to the dining area. The food dispenser has two stacked plates, each with a capacity of 24 food items (**b**). A water port in the waiting area allows mice to drink and thus consume more food. Two pairs of infrared (IR) break-beam sensors detect a mouse moving from the waiting to the dining area. A door is used to block access to the dining area during food delivery. Three cameras record mouse behavior and a microphone records bite sound. Mouse drawings in **a** were adapted from scidraw.io (https://scidraw.io/drawing/122 and https://scidraw.io/drawing/96). **a.** Behavioral sequence in the Mouse Restaurant and signals used for task control. Behavioral stages in red were recorded in the dining area. **b.** Processing of audio signal for bite detection. Audio signal was band-pass filtered (800-8,000 Hz), rectified, smoothed (5-ms Gaussian window), and thresholded (4 × s.d. above mean) to detect bite events (red circles). Blue rectangle indicates the time window enlarged on the right. **e-g.** Image sequences showing manually labeled action motifs observed in angel-hair pasta eating. Images in each panel represent the start (left), middle, and end (right) of each action. Arrows in **e** highlight the jaw as it opens to retrieve the pasta. Arrows in **f** hightlight the tongue as it brings the pasta into the mouth. Arrows in **g** indicate the upward body movement leading to the sitting posture. **h.** An example image taken at the start of a hand-withdraw event, in which a mouse raises its hands toward the mouth (arrow) to start a handle-eat bout following a chewing phase. Right panel shows Z-axis trajectory of the right hand before and after the hand-withdraw event, with the blue line indicating the time of the hand withdraw shown in the left image. **i.** Image sequences showing unimanual (upper sequence) and bimanual (lower sequence) adjustments through release and re-grasp movements to reposition the hands on the pasta. Arrows in **i** highlight the release and re-grasp hand movements.

**Extended Data Fig. 5.**
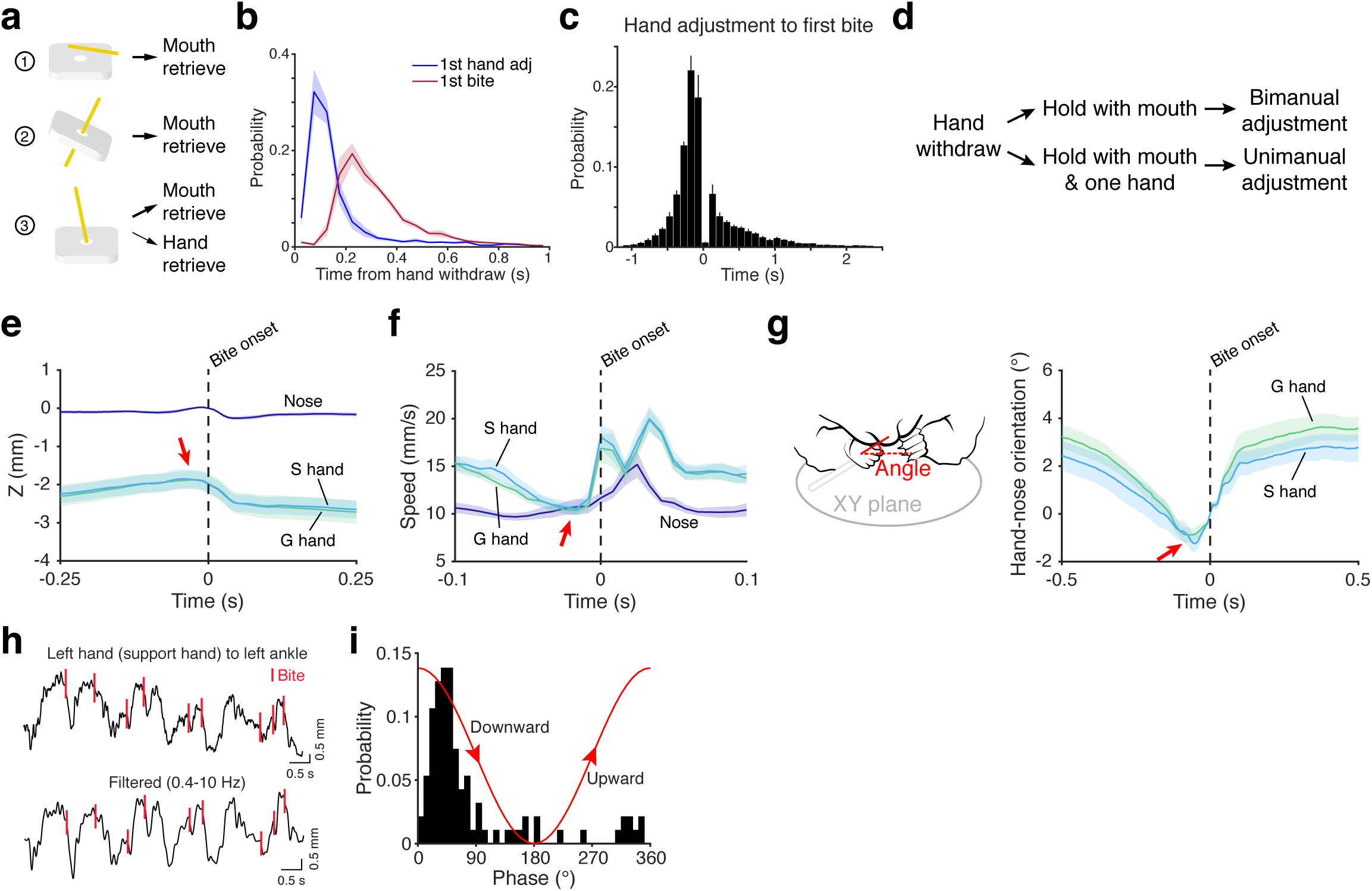
Hand adjustment and pasta bite both involve oromanual coordination. Related to Fig. 2. **a.** Configurations of a piece of 15-mm angel-hair pasta when delivered to the dining area. 3D-printed holders were used to load the pasta into the food dispenser in **Extended Data Fig. 4b**. For configuration 3, mice occasionally retrieved the pasta with the hands instead of the mouth. Trials with hand retrieval were not included in the analysis due to low occurrence. **b.** Probability distribution of the time of the first hand adjustment and the first bite in each handle-eat bout (n = 7 mice). Shades around mean denote ± s.e.m. **c.** Probability distribution of the time from hand adjustments to the first bite in each handle-eat bout. The proportion of hand adjustments made before the first bite for 7 mice is 69.2 ± 2.2 %, indicating hand adjustment mainly occurs before the first bite in each bout. Data are mean ± s.e.m. **d.** Action sequences for unimanual and bimanual adjustments. **e, f.** Average Z-axis position (**e**) and speed (**f**) of nose and hands aligned to bite onset showing that pasta biting involves coordinated bimanual and jaw movement. Note that the downward hand movement starts before the bite (arrows in **e, f**). The two peaks in **f** are likely due to the breaking of pasta. **g.** Average hand-nose orientation aligned to bite onset, showing a downward hand movement relative to the nose (mouth) before a bite (red arrow) to snap the pasta. The schematic depicts the angle of hand-nose orientation (left panel). **h.** The relationship between up-down hand movements and bite, shown as the Z-axis left-hand trajectory overlaid with bite events. Bite mostly occurs during the downward swing of the hand to snap the pasta. Left-ankle position was used as a reference to compute the trajectory, which was then band-pass filtered (0.4 - 10 Hz, lower panel) to compute the hand-movement phase. **i.** Probability distribution of the phases of support-hand movement at the time of bites from an example trial. The red cosine curve indicates hand movement. Shades around mean denote ± s.e.m in **e-g** for 9 mice.

**Extended Data Fig. 6.**
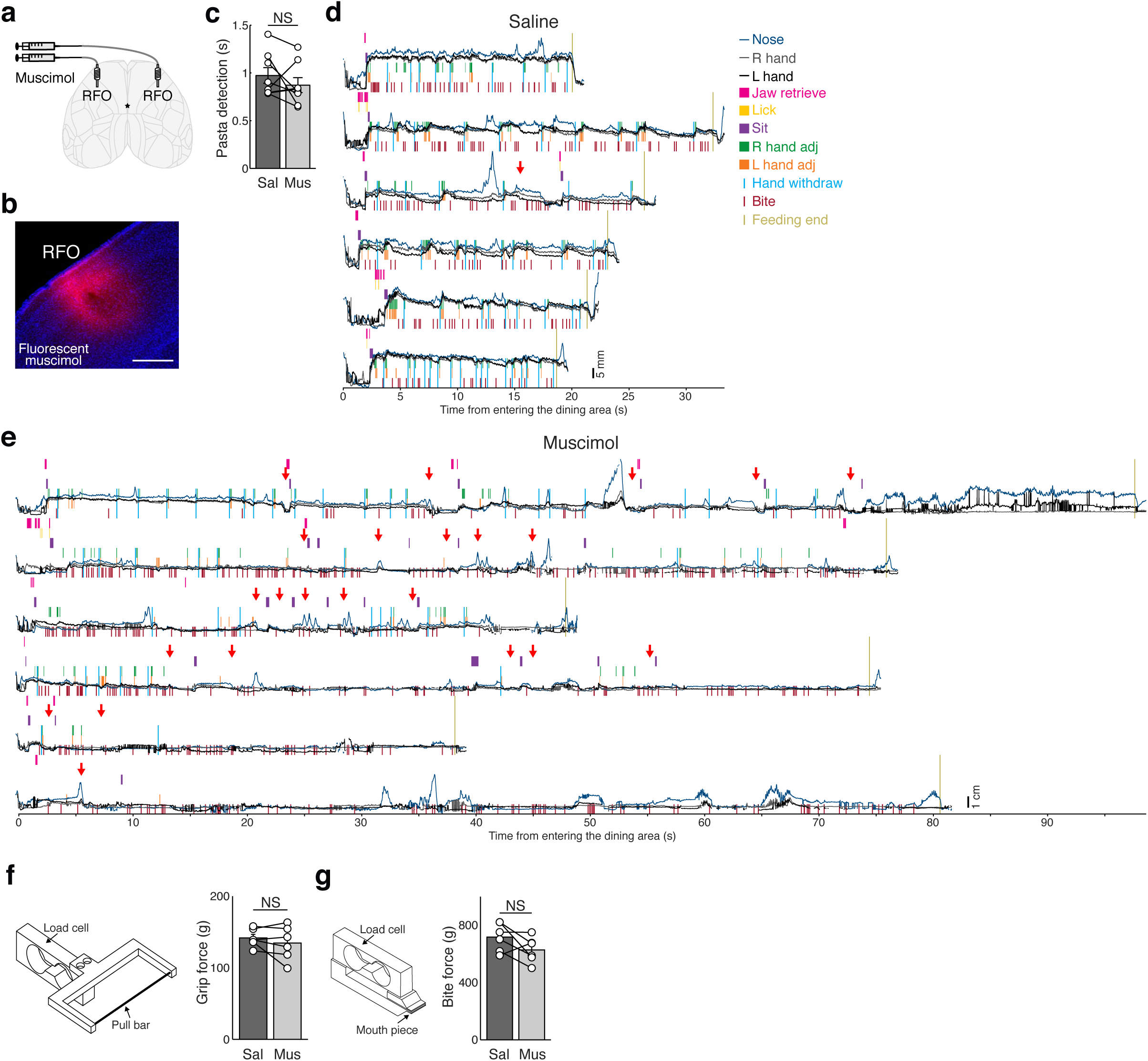
Muscimol inhibition in RFO impairs hand recruitment in pasta eating but not bite and grip force. Related to Fig. 2. **a.** Schematic of bilateral muscimol infusion into the RFO. **b.** Representative diffusion pattern of BODIPY-tagged muscimol (red; 1 µl) in the RFO of coronally sectioned (75 µm) tissue stained with DAPI (blue). Scale bar, 500 µm. **c.** Muscimol inhibition did not impair pasta detection (n = 8 mice). **d, e.** Actograms of example trials of a mouse following bilateral saline (**d**) or muscimol (**e**) infusion. In muscimol trials, the mouse usually did not adopt a sitting posture (nose and hand position close to the ground), bit the pasta on the ground without recruiting hands, and often dropped the pasta (red arrows) during eating. In muscimol trials, feeding time was significantly longer. Sometimes, a mouse left the dining area without finishing its pasta, or the pasta flew out of the dining area after a bite due to uncoordinated oromanual movements. **f, g**. Muscimol inhibition did not impair grip force (**f**) or bite force (**g**). n = 6 mice. Schematics in **f** and **g** depict the apparatus used to measure grip force and bite force respectively. Data are mean ± s.e.m in **c**, **f, g**. NS, not significant, two-sided paired t-test in **c**, **f, g**.

**Extended Data Fig. 7.**
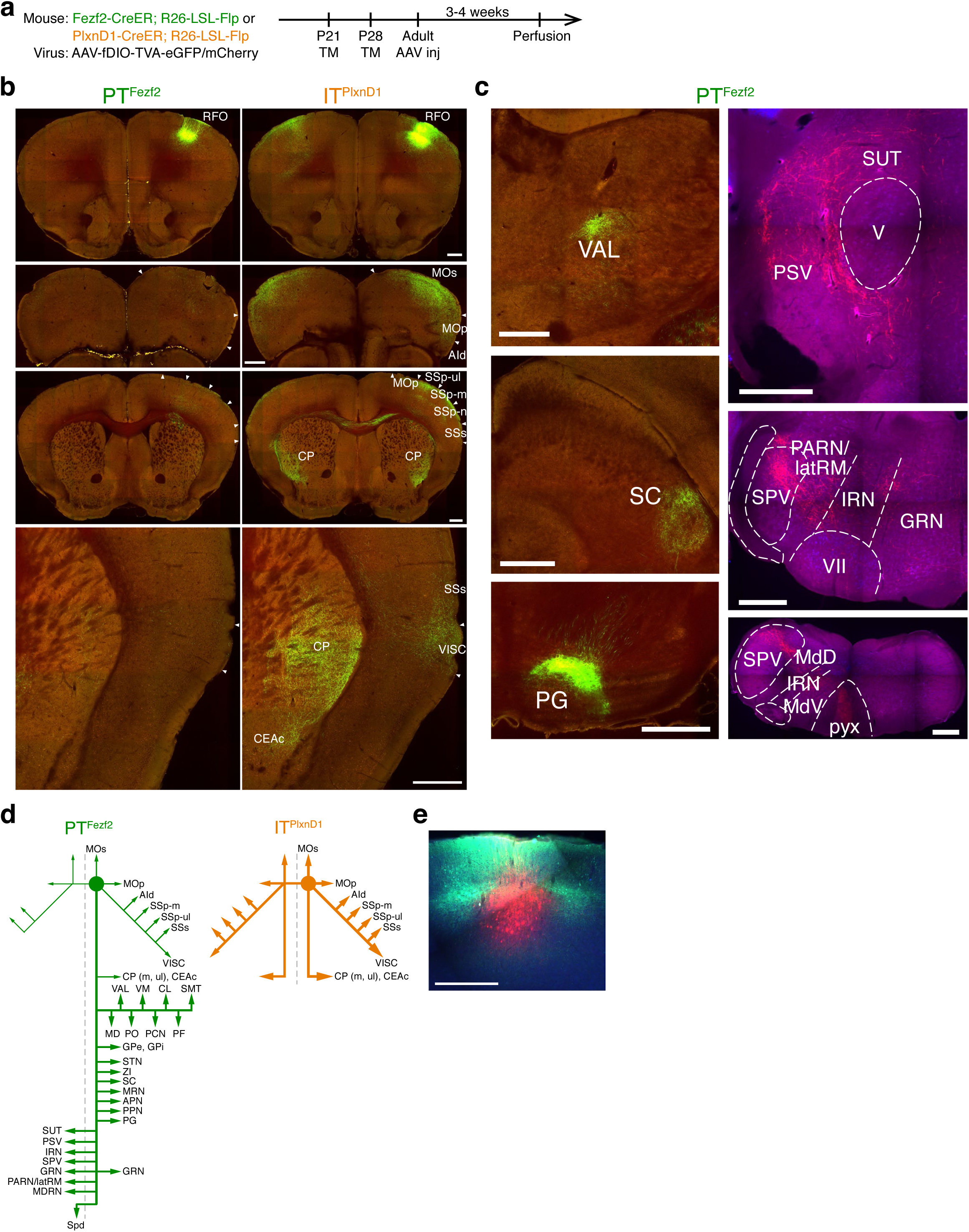
Brain-wide projection targets of PTs^Fezf2^ and ITs^PlxnD1^ in RFO. Related to Fig. 3. **a.** Strategy and timeline for anterograde tracing of PTs^Fezf2^ and ITs^PlxnD1^ in the RFO. TM, tamoxifen. **b.** Images at the RFO injection site (first row) and selected projection targets: eGFP expression from Flp-activated viral vector (green) and background autofluorescence (red). PTs^Fezf2^ show a weak projection to the cortex and striatum whereas ITs^PlxnD1^ show a strong bilateral projection to the cortex and striatum. **c.** Images of selected subcortical projection targets of PTs^Fezf2^. Left panels show eGFP expression from Flp-activated viral vector (green) and background autofluorescence (red). Right panels show mCherry expression from Flp-activated viral vector (red) and Nissl staining (blue). PT^Fezf2^ axons form the pyramidal decussation and enter the spinal cord (bottom right panel). **d.** Schematic depicting main efferent targets of RFO PTs^Fezf2^ and ITs^PlxnD1^. ITs^PlxnD1^ project bilaterally to multiple cortical areas, the ventrolateral striatum, and CEAc. PTs^Fezf2^ project weakly within the cerebral cortex and striatum but strongly to subcortical structures at all levels. **e.** Image at RFO injection site from simultaneous PTs^Fezf2^ and ITs^PlxnD1^ tracing. PTs^Fezf2^, red; ITs^PlxnD1^, green. Scale bar, 500 µm. AId, agranular insular area, dorsal part; APN, anterior pretectal nucleus; CEAc, central amygdalar nucleus, capsular part; CL, central lateral nucleus of the thalamus; CP, caudoputamen; GPe, globus pallidus, external segment; GPi, globus pallidus, internal segment; GRN, gigantocellular reticular nucleus; IRN, intermediate reticular nucleus; latRM, lateral rostral medulla; m, mouth; MD, mediodorsal nucleus of the thalamus; MdD, medullary reticular nucleus, dorsal part; MDRN, medullary reticular nucleus; MdV, medullary reticular nucleus, ventral part; MOp, primary motor area; MOs, secondary motor area; MRN, midbrain reticular nucleus; PARN, parvicellular reticular nucleus; PCN, paracentral nucleus; PF, parafascicular nucleus; PG, pontine gray; PO, posterior complex of the thalamus; PPN, pedunculopontine nucleus; PSV, principal sensory nucleus of the trigeminal; pyx, pyramidal decussation; SC, superior colliculus; SMT, submedial nucleus of the thalamus; Spd, spinal cord; SPV, spinal nucleus of the trigeminal; SSp-m, primary somatosensory area, mouth; SSp-n, primary somatosensory area, nose; SSp-ul, primary somatosensory area, upper limb; SSs, secondary somatosensory area; STN, subthalamic nucleus; SUT, supratrigeminal nucleus; ul, upper limb; V, motor nucleus of trigeminal; VAL, ventral anterior-lateral complex of the thalamus; VII, facial motor nucleus; VISC, visceral area; VM, ventral medial nucleus of the thalamus; ZI, zona incerta.

**Extended Data Fig. 8.**
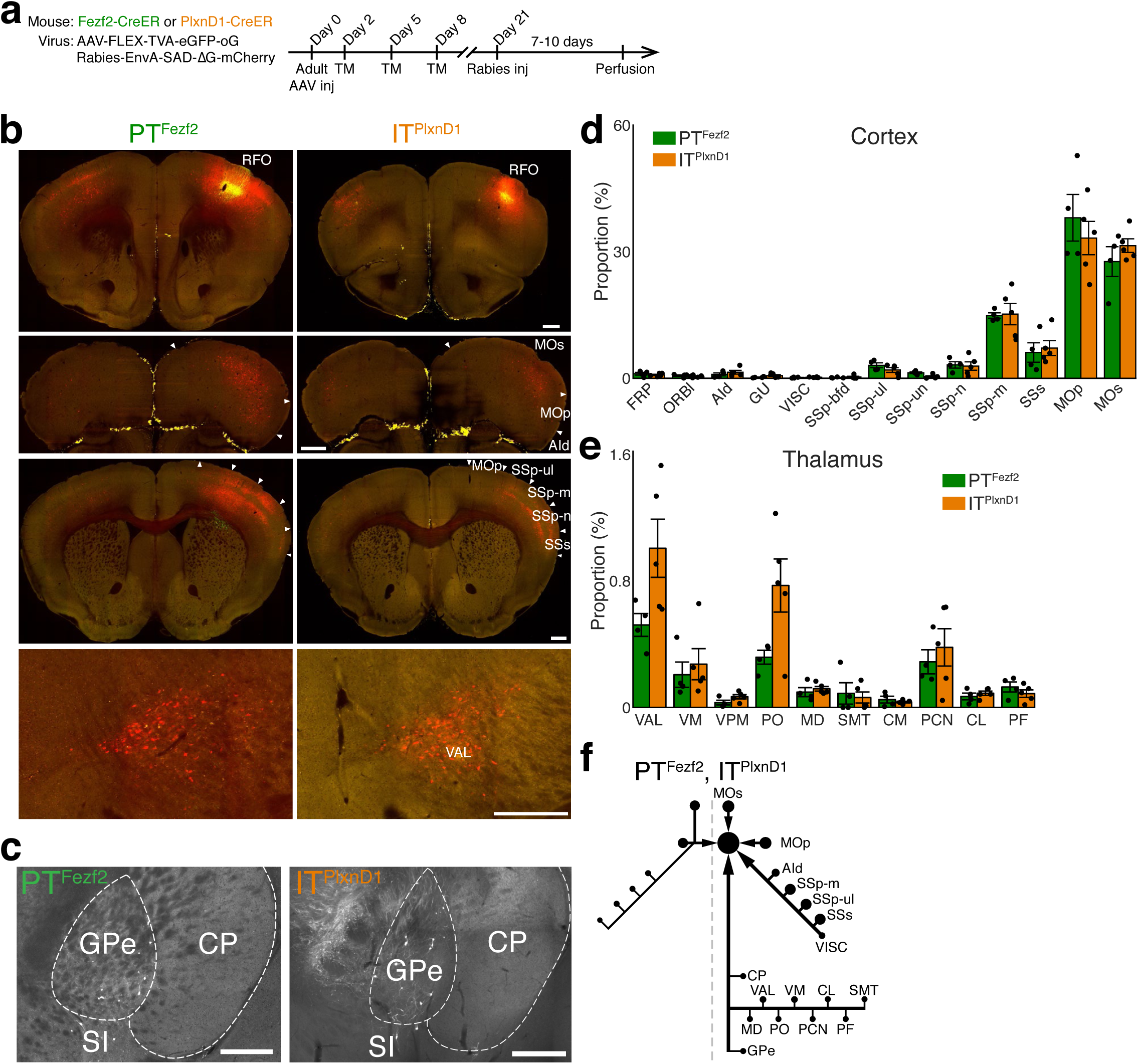
Brain-wide monosynaptic inputs to PTs^Fezf2^ and ITs^PlxnD1^ in RFO. Related to Fig. 3. **a.** Strategy and timeline for retrograde monosynaptic rabies tracing of PTs^Fezf2^ and ITs^PlxnD1^ in the RFO. TM, tamoxifen. **b.** Images at RFO injection site (first row) and selected afferent sources: mCherry expression from rabies viral vector (red) and eGFP expression from Cre-activated starter virus (green). Both PTs^Fezf2^ and ITs^PlxnD1^ receive afferents from cortical areas and the thalamus. **c.** Images showing input cells in the GPe that monosynaptically connect to PTs^Fezf2^ (left panel) and ITs^PlxnD1^ (right panel) in the RFO. **d, e.** Proportion of input cells in cortical areas and thalamic nuclei (4 *Fezf2* and 5 *PlxnD1* mice). Data are mean ± s.e.m. **f.** Schematic depicting input sources to PTs^Fezf2^ and ITs^PlxnD1^ in the RFO from cortical areas, the thalamus, and GPe. The size of the nodes reflects the number of input cells. Scale bar, 500 µm in **b** and **c**. AId, agranular insular area, dorsal part; CL, central lateral nucleus of the thalamus; CM, central medial nucleus of the thalamus; CP, caudoputamen; FRP, frontal pole; GPe, globus pallidus, external segment; GU, gustatory areas; MD, mediodorsal nucleus of the thalamus; MOp, primary motor area; MOs, secondary motor area; ORBl, orbital area, lateral part; PCN, paracentral nucleus; PF, parafascicular nucleus; PO, posterior complex of the thalamus; SI, substantia innominata; SMT, submedial nucleus of the thalamus; SSp-bfd, primary somatosensory area, barrel field; SSp-m, primary somatosensory area, mouth; SSp-n, primary somatosensory area, nose; SSp-ul, primary somatosensory area, upper limb; SSp-un, primary somatosensory area, unassigned; SSs, secondary somatosensory area; VAL, ventral anterior-lateral complex of the thalamus; VISC, visceral area; VM, ventral medial nucleus of the thalamus; VPM, ventral posteromedial nucleus of the thalamus.

**Extended Data Fig. 9.**
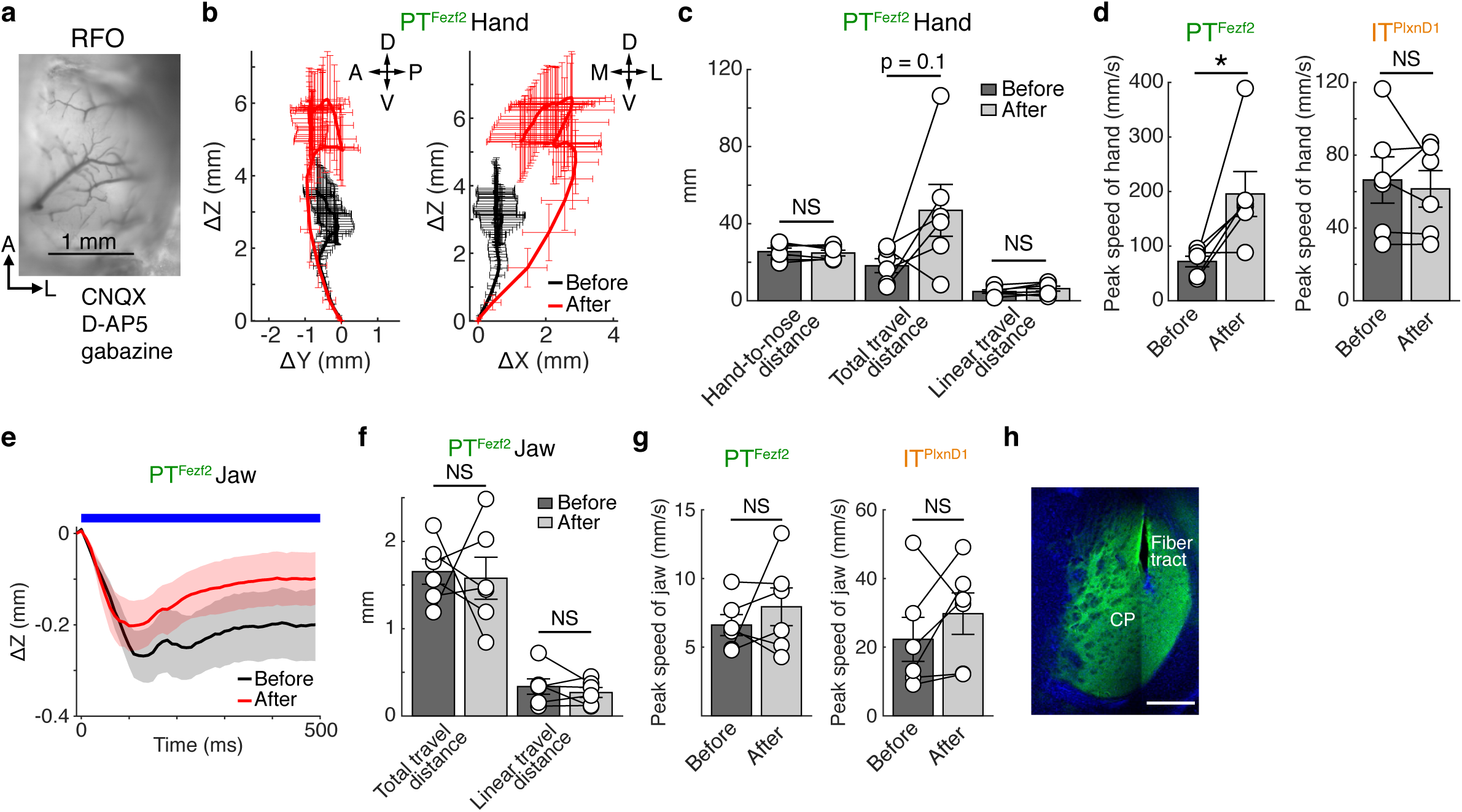
Parallel PT^Fezf2^-corticofugal and IT^PlxnD1^-striatal pathways convey their activation effects on forelimb and mouth movements. Related to Fig. 3. **a.** Image of local cortical application of CNQX, D-AP5, gabazine in the RFO (see Methods). **b.** 2D projections of left-hand trajectories following stimulation of RFO PTs^Fezf2^. Black and red curves represent trajectories before and after drug application, respectively. Movement directions are indicated as: A, anterior; P, posterior; D, dorsal; V, ventral; M, medial; L, lateral. **c.** Hand-to-nose distance, total- and linear-travel distance of hand movement before and after drug application. **d.** Peak speed of hand movement before and after drug application. **e.** Movement trajectory of the jaw following stimulation of RFO PTs^Fezf2^. Black and red curves represent trajectories before and after drug application, respectively. Blue bar represents stimulation window. Shading around mean denotes ± s.e.m. **f.** Total- and linear-travel distance of jaw movement before and after drug application. **g.** Peak speed of jaw movement before and after drug application. **h.** Image of a coronal section showing IT^PlxnD1^ axons and fiber tract in the striatum. Scale bar, 0.5 mm. n = 6 mice for PTs^Fezf2^ or ITs^PlxnD1^ in **b**-**g**. Movement trajectories were normalized to the start position in **b**, **e**. Data are mean ± s.e.m in **b**-**d**, **f**, and **g**. *p < 0.05, two-sided paired t-test; NS, not significant.

**Extended Data Fig. 10.**
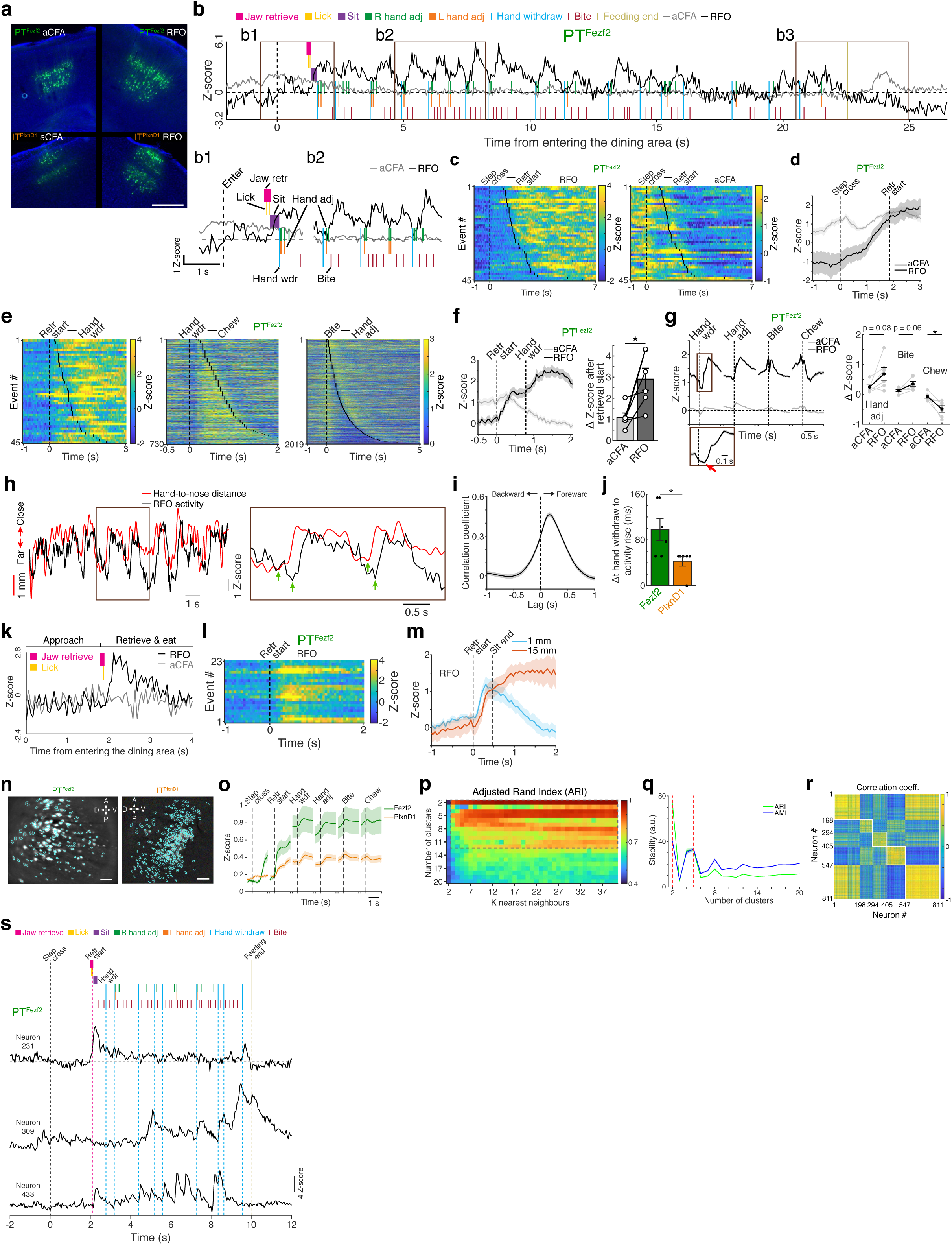
PT^Fezf2^ and IT^PlxnD1^ activity in RFO correlate with oromanual manipulation. Related to Fig. 4. **a.** Coronal sections showing PTs^Fezf2^ and ITs^PlxnD1^ in the RFO and aCFA expressing GCaMP7f from AAV infection. Scale bar, 500 µm. **b.** Single-trial calcium activity traces of PTs^Fezf2^ in the RFO (black) and aCFA (gray) of a mouse eating 15-mm angel-hair pasta overlaid on the actogram. Example time windows are highlighted (**b1-b3**) and two of them are expanded (**b1** and **b2**). The rise of aCFA activity at time 0 (dashed line) correlates with crossing a step when entering the dining area (see Fig. 2a). **c.** Heat maps of PT^Fezf2^ population activity in the RFO and aCFA aligned to entry into the dining area and sorted by the start of pasta retrieval. **d.** Average population activity of PTs^Fezf2^ in the RFO and aCFA aligned to entry into the dining area (n = 6 mice). Vertical dashed lines indicate the average time to the retrieval start. **e.** Heat maps of RFO PT^Fezf2^ population activity aligned to retrieval start (left), hand withdraw (middle), and bite (right). Activity traces were sorted by the earliest hand withdraw (left), chew (middle), and hand adjustment (right) events, respectively. **f, g.** Average PT^Fezf2^ population activity in the RFO and aCFA aligned to retrieval start (**f**; left panel) and hand withdraw, hand adjustment, bite, and chew (**g**; left panel). Vertical dashed lines indicate average time to the first hand withdraw in **f**. Changes in population activity are shown in the right panels. RFO PT^Fezf2^ activity rise after the onset of hand withdraw with a lag (red arrow in the expanded window of **g)**. n = 6 mice; *p < 0.05, two-sided paired t-test. **h.** Correlation between RFO IT^PlxnD1^ population activity and hand-to-nose distance. Boxed time window is expanded on the right. Green arrows indicate the onset of signal rise. **i.** Averaged correlation coefficient of RFO IT^PlxnD1^ population activity with hand-to-nose distance shifted in time from an example mouse. **j.** Time from the onset of hand withdraw to the rise of population activity (n = 6 mice for PTs^Fezf2^ or ITs^PlxnD1^; *p < 0.05, two-sided Wilcoxon rank-sum test). The rise of IT^PlxnD1^ activity preceded that of PT^Fezf2^ activity. **k, l.** Single-trial PT^Fezf2^ calcium activity in the RFO and aCFA as a mouse consumed 1-mm angel-hair pasta without sitting up and hand recruitment (**k**). Key actions (colored annotations) were overlaid on the activity traces. Corresponding calcium activity (**l**) were aligned to retrieval start for 1-mm pasta. **m.** Averaged RFO PT^Fezf2^ population activity aligned to retrieval start for 15-mm and 1-mm angel-hair pasta (n = 7 mice). Vertical dashed lines indicate the average time to establish a sitting posture when eating 15-mm pasta. Activity remained high when mice handled and ate 15-mm pasta but declined when eating 1-mm pasta without sitting and hand recruitment. **n.** Extracted PT^Fezf2^ and IT^PlxnD1^ ROIs overlaid on the maximum projection of miniscope calcium imaging video. A, anterior; P, posterior; D, dorsal; V, ventral. Scale bar: 100 µm. **o.** Average activity of all recorded neurons aligned to different action motifs. **p.** Heat map of ARI (adjusted rand index), mean of 100 bootstrap sessions, for different combinations of cluster numbers and K nearest neighbours. Dashed rectangle region indicates the hyperparameter space with high clustering stability. **q.** Variance-adjusted ARI and AMI (adjusted mutual information), mean across bootstraps/SD, for different number of clusters. Shading around mean denotes ± s.e.m across k-NN from 2 to 40. Red dashed lines, two most stable clusterizations. **r.** Correlation between average activity aligned to different action motifs of neurons assigned to the five clusters. White squares depict neurons in the same functional cluster. **s.** Activity of three simultaneously recorded PTs^Fezf2^ in an exemplar trial. The actogram is shown at the top. Horizontal dashed lines depict Z-score of 0. The two-cluster approach (**q**) only broadly divided the neurons into two groups either with general increased or decreased activity to all action motifs (data not shown), which likely underestimated the heterogeneity of neuronal activity patterns in the RFO. We thus focused on the scheme of five clusters that showed high activity correlation for neurons in the same clusters (**r**). Shading around mean denotes ± s.e.m in **d, f, g**, **i**, **m**, **o**, **q**. Data are mean ± s.e.m in **f, g, j**.

**Extended Data Fig. 11.**
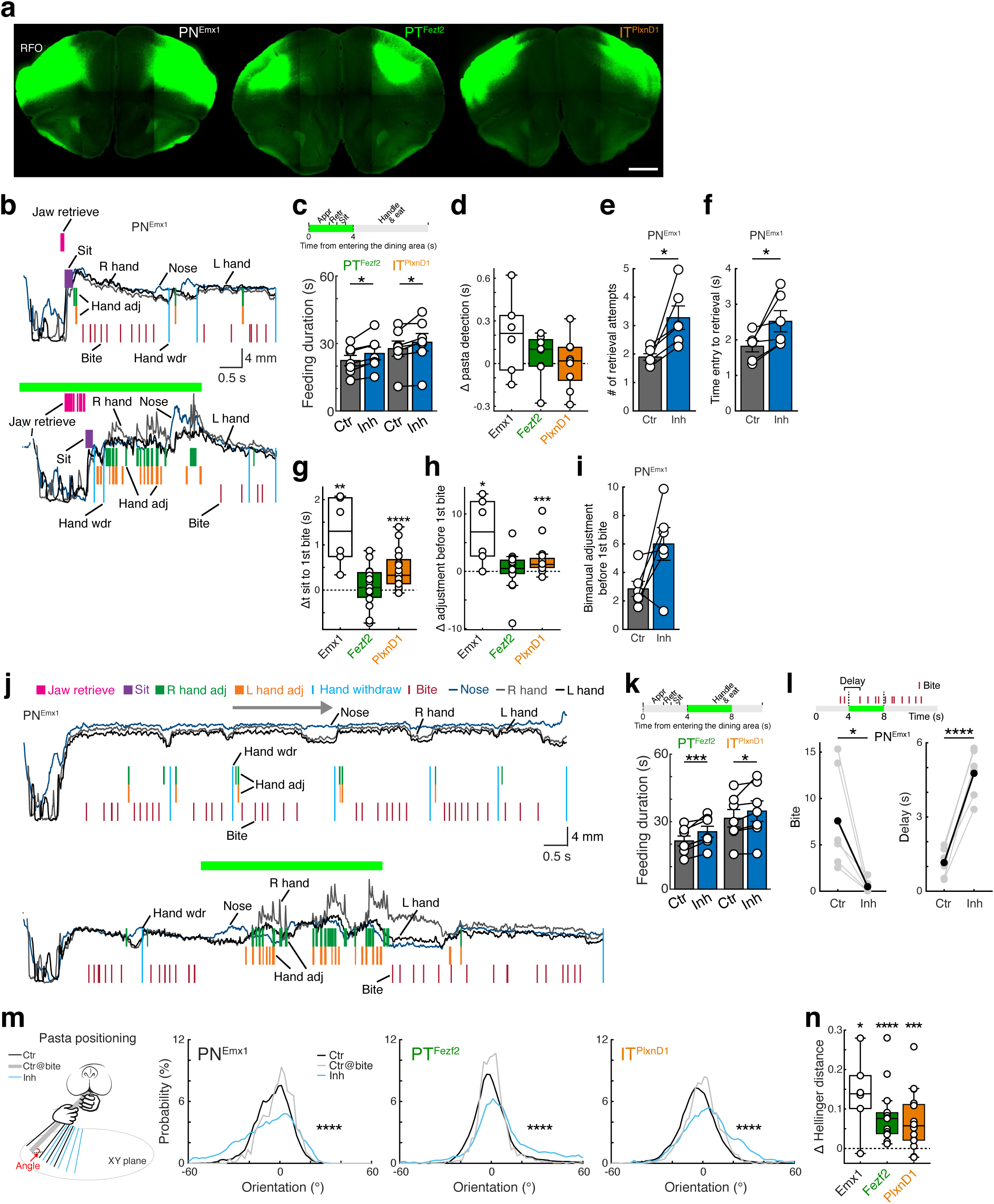
PTs^Fezf2^ and ITs^PlxnD1^ in RFO contribute to oromanual manipulation. Related to Fig. 5. **a.** Coronal sections showing PNs^Emx1^, PTs^Fezf2^, and ITs^PlxnD1^ in the RFO that were infected by Cre-dependent AAV-DIO-GtACR1-eYFP injection in the corresponding driver mouse. Scale bar, 1 mm. **b.** Actograms of a mouse in control (upper) and PN^Emx1^ inhibition (lower; green bar) trials. Z-axis trajectories of nose, right and left hands are shown. Actions are colored and labeled. **c.** Optogenetic inhibition of RFO PTs^Fezf2^ or ITs^PlxnD1^ in the retrieve-eat stage increased feeding duration. Schematic at the top depicts the inhibition scheme. Green bar indicates 4-s inhibition. **d.** PN inhibition did not impact the time taken for pasta detection compared to control trials. **e, f.** PN^Emx1^ inhibition interfered with pasta retrieval, measured as increased number of retrieval attempts (i.e., retrieval jaw movements) (**e**) and lengthened the time from entry to retrieval (**f**). **g.** PN^Emx1^ and IT^PlxnD1^ inhibition delayed the first bite after adopting a sitting posture. **h.** PN^Emx1^ and IT^PlxnD1^ inhibition increased total number of hand adjustment before the first bite. **i.** Changes in the number of bimanual adjustment before the first bite following PN^Emx1^ inhibition. **j.** Actograms of a mouse at the handle-eat stage in control (top) and PN^Emx1^ inhibition (bottom) trials. Z-axis trajectories of nose and two hands are shown. PN^Emx1^ inhibition led to substantially increased hand adjustments but no bite. The top horizontal arrow indicates a control handle-eat bout. **k.** Optogenetic inhibition of RFO PTs^Fezf2^ or ITs^PlxnD1^ in the handle-eat stage increased feeding duration. Schematic at the top depicts the inhibition scheme. Green bar indicates 4-s inhibition. **l.** PN^Emx1^ inhibition resulted in decreased number (left) and increased delay (right) of bites. Red ticks in the top schematic indicate bite events. **m.** Probability distributions of pasta orientation during the handle-eat stage in control and PN inhibition trials; gray traces denote probability distributions at the time of bite/snap in control trials. Schematic shows exemplar pasta orientation for different conditions. XY plane denotes the ground plane. Orientation was normalized for each mouse based on the average bite orientation of control trials and pooled across mice (****p < 0.001, Control vs Inhibition, Kolmogorov-Smirnov test). **n.** Probability distribution of all pasta orientations in control trials was more similar to the probability distribution of bite orientations in control trials compared with that of all pasta orientations in inhibition trials, quantified as difference in Hellinger distance. The smaller the Hellinger distance, the more similar the two probability distributions. n = 7 mice for PTs^Fezf2^ or ITs^PlxnD1^ in **c, k**. n = 8 mice for PTs^Fezf2^ and 9 mice for ITs^PlxnD1^ in **d**. n = 15 mice for PTs^Fezf2^ and 16 mice for ITs^PlxnD1^ in **g, h, m, n**. n = 6 mice for PNs^Emx1^ in **d-i** and **l-n**. *p < 0.05, **p < 0.01, ***p < 0.005, ****p < 0.001, two-sided paired t-test and two-sided Wilcoxon signed-rank test in **c-i**, **k, l, n**.

**Extended Data Fig. 12.**
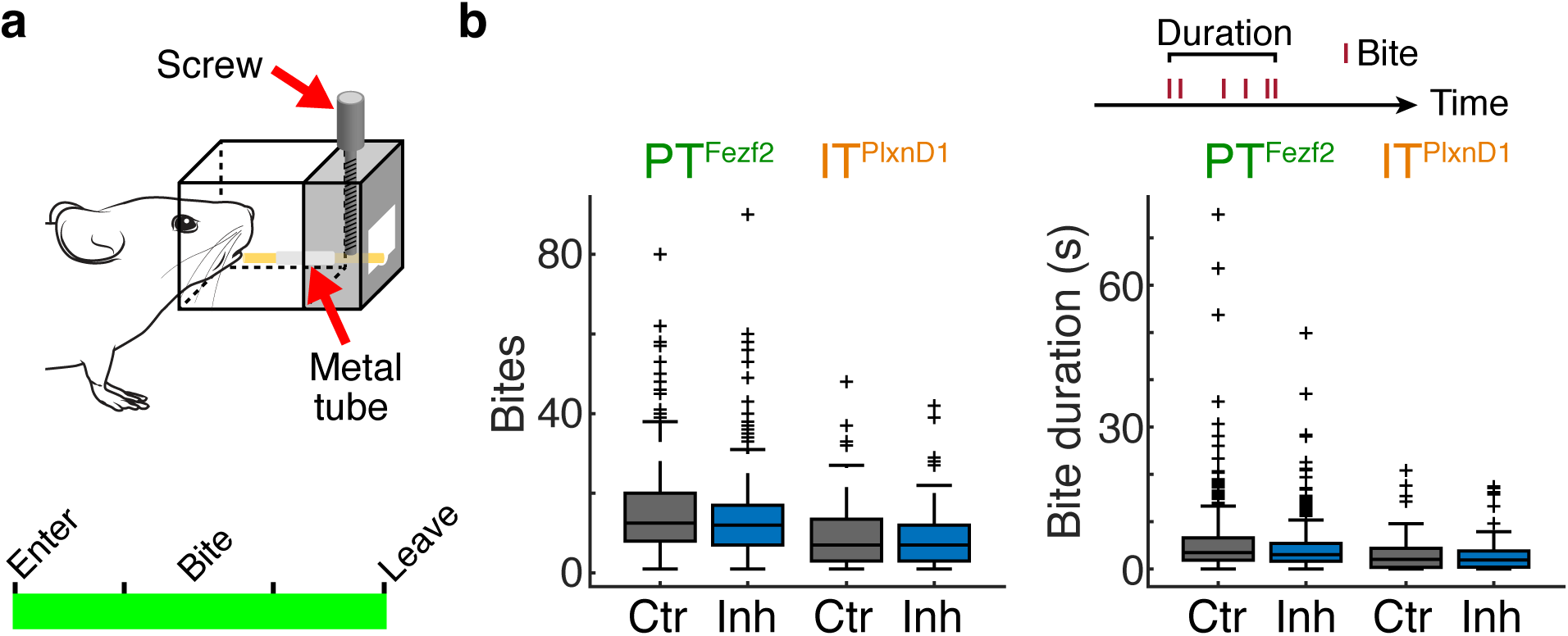
Inhibiting PTs^Fezf2^ or ITs^PlxnD1^ in RFO does not impair bite using mouth. Related to Fig. 5. **a.** Schematic of pasta-bite apparatus. Angel-hair pasta is inserted into a metal tube and secured in place with a screw. A small segment (∼ 3 mm) of the pasta projects from the tube allowing a mouse to bite off the pasta segment without hand use. Bottom panel shows inhibition scheme, which covers the whole trial period (green bar). **b.** Number of bites (left panel) and duration taken (right panel) to bite a pasta segment. Red ticks in the top schematic indicate bite events. n = 7 mice for PTs^Fezf2^ and 3 mice for ITs^PlxnD1^. The mouse drawing in **a** was adapted from scidraw.io (https://scidraw.io/drawing/94).

## SUPPLEMENTARY VIDEOS

**Supplementary Video 1. Optogenetic activation of PTs^Fezf2^ in different locations of the dorsal cortex of head-fixed mouse.** Optogenetic activation of PTs^Fezf2^ in posterior Caudal Forelimb Areas (pCFA: P 1.125, L 1.125; 0.5 s) in a head-fixed mouse induces a lateral abduction of the left forelimb, with digit opening and extension and elbow extension. Associated facial movements include vibrissae whisking and eyelid opening. Optogenetic activation of PTs^Fezf2^ in medial Caudal Forelimb Area (mCFA: A 0, L 1.5; 0.5 s) in a head-fixed mouse induces treading (up/down) movements of the left forelimb. With stimulation onset the forelimb is raised by elbow flexion and then lowered by elbow extension (repeated a number of times). Digit flexion follows elbow flexion and digit extension leads elbow extension. Vibrissae whisk with a similar rhythm to the treading movement. The movement has features of a placing response in which a hand attempts to contact and obtain support from a surface. Optogenetic activation of PTs^Fezf2^ in anterior Caudal Forelimb Area (aCFA: 0.75, L 1.875; 0.5 s) in a head-fixed mouse induces a stepping or reaching-like forelimb movement. The upward movement involves sequentially, elbow, wrist, and digit flexion followed by extension. At the apex of the movement the limb is in a relaxed posture. Eyelid opening and whisking accompany the movement. The movement has features resembling reaching or stepping. Optogenetic activation of PTs^Fezf2^ in Rostral Forelimb Orofacial area (RFO: A 1.5, L 2.25; 0.5 s) in a head-fixed mouse induces left hand adduction to the body midline with hand supination and digit flexing and closing. Associated facial movement includes two cycles of jaw opening and closing with lateral leftward tongue protrusion.

**Supplementary Video 2. Optogenetic activation of ITs^PlxnD1^ in Rostral Forelimb Orofacial area of head-fixed mouse.** Optogenetic activation (ITs^PlxnD1^ in RFO: A 1.5, L 2.25; 0.5 s) in a head-fixed mouse induces bilateral digit flexion and closing followed by elbow flexion and adduction of both hands toward the body midline. Adduction and flexion at the shoulders then raise both hands to the mouth. The movement has features of eating or grooming.

**Supplementary Video 3. Optogenetic activation of PTs^Fezf2^ or ITs^PlxnD1^ in Rostral Forelimb Orofacial area of free-moving mouse.** Optogenetic activation of PTs^Fezf2^ in RFO (A 1.7, L 2.5; 0.5 s) in a free-moving mouse induces shoulder adduction that carries the left hand, with associated hand supination, toward the body midline. Ipsiversive head turning and lowering bring the snout close to the hand. The right hand maintains body postural support. Optogenetic activation of ITs^PlxnD1^ in RFO (A 2, L 2.625; 0.5 s) in a free-moving mouse interrupts ongoing behavior. The mouse adopts a sitting posture and concomitant bilateral shoulder adduction brings both hands to the body midline. During adduction, the digits flex and close and contact the mouth. At stimulation termination, the hands are replaced on the floor.

**Supplementary Video 4. A dorsal view of the Mouse Restaurant**. The mouse leaves the waiting area, proceeds down a corridor and steps down a small step to enter the dining area to find and eat a food pellet. The mouse’s movements are enabled by opening the “door” to allow access to the dining area and by positioning a “table”, containing a food item e.g., food pellet, angel-hair pasta, in the dining area.

**Supplementary Video 5. Angel-hair pasta eating after RFO saline or muscimol infusion.** After saline infusion (Rostral Forelimb Orofacial area, RFO), a mouse enters the dining area and finds a 15mm piece of angel-hair pasta. It sniffs and whisks the pasta and then directs its snout to an end of the pasta where with tongue/mouth movements it grasps the pasta with the incisors. Pasta positioning in the mouth induces the adoption of a sitting posture on the haunches and concurrent raising of both hands to grasp the pasta. Bilateral hand adjustments with assistance of the mouth position the pasta in the mouth in an oblique orientation for biting. The pasta is consumed by repeated acts of positioning, biting, and chewing mediated by coordinated oromanual movements. The tracking of different body parts and the pasta are shown. After muscimol infusion Mouse 1 identifies the pasta by sniffing. It is clumsy in picking up the pasta by mouth, does not seek out the end of the pasta for mouth purchase, does not use its tongue/mouth to grasp the pasta and makes little use of its hands for food retrieval from the mouth or pasta manipulation. The pasta is consumed from the floor mainly using mouth movements. After muscimol infusion Mouse 2 identifies the pasta by sniffing, does not seek out the end of the pasta for tongue/mouth purchase, and picks it up in the middle with its mouth. It lifts the hands to grasp the pasta but fails to manipulate the pasta or remove it from its mouth to reorient it into a position for biting. The mouse ends up breaking the pasta in half.

**Supplementary Video 6**. **Input-output tracing of RFO PTs^Fezf2^ and ITs^PlxnD1^.** Whole-brain stacked images of PT^Fezf2^ projection show axons mainly in the ventrolateral part of the ipsilateral striatum, thalamus (e.g., VAL, PO, and PF), lateral superior colliculus, pons, and medulla. The axons are mainly in the contralateral medulla and eventually crossed the midline at the pyramidal decussation to innervate the spinal cord. Whole-brain stacked images of IT^PlxnD1^ projection show axons in cortical areas (e.g., MOs, MOp, SSp-ul, SSp-m, SSp-n, SSs, AId, and VISC) and the ventrolateral part of the striatum of both hemispheres. In addition, ITs^PlxnD1^ project bilaterally to the capsular part of the central amygdala nucleus. Whole-brain stacked images of PT^Fezf2^ or IT^PlxnD1^ input show rabies-virus labeled input cells mainly from cortical areas (e.g., MOs, MOp, SSp-ul, SSp-m, SSp-n, SSs, AId, and VISC) and the thalamus (e.g., VAL, PO, PCN, and VM). VAL, ventral anterior-lateral complex of the thalamus; PO, posterior complex of the thalamus; PF, parafascicular nucleus; RFO, Rostral Forelimb Orofacial area; PT, pyramidal tract; MOs, secondary motor area; MOp, primary motor area; AId, agranular insular area, dorsal part; SSp-ul, primary somatosensory area, upper limb; SSp-m, primary somatosensory area, mouth; SSp-n, primary somatosensory area, nose; SSs, secondary somatosensory area; VISC, visceral area; IT, intratelencephalic. PCN, paracentral nucleus; VM, ventral medial nucleus of the thalamus.

**Supplementary Video 7. Retrieve-eat stage of pasta eating in control or optogenetic inhibition trials.** In control trial, a mouse grasps angel-hair pasta (15 mm) by orienting its head so that it can grasp the end of the pasta. The mouse then immediately adopts a sitting posture, uses its hands to take the pasta to help orient the pasta in its mouth. Using oromanual manipulation, it proceeds to bite pieces from the pasta. Optogenetic inhibition of RFO (Rostral Forelimb Orofacial area) PNs^Emx1^ (4 sec duration top left; 15mm-angel hair pasta) starts with mouse entry to the dining area. The mouse does not orient the mouth to the end of the pasta and grasps the pasta with its mouth after the 5^th^ attempt. It then immediately adopts a sitting posture and grasps the pasta with its hands, but does not orient its mouth to the end of the pasta but bites the pasta in its middle. Optogenetic inhibition of RFO PTs^Fezf2^ (4 sec duration top left; 15mm-angel hair pasta) begins as the mouse enters the dining area. The mouse orients its mouth to the end of the pasta but only grasps the pasta after the 6^th^ attempt. Once the pasta is grasped, the mouse immediately adopts a sitting posture and orients its mouth to the end of pasta to bite.

**Supplementary Video 8. Pasta-bite test with or without RFO PN-type inhibition.** In the control trial of the pasta-bite test, a mouse approaches, detects, orients its mouth, and successfully bites a piece of angel-hair pasta that projects horizontally from a holder located in the aperture. Optogenetic inhibition (PTs^Fezf2^; whole trial) in Rostral Forelimb Orofacial area does not affect approach, detection, head orient, and successful bite of a piece of angel-hair pasta that projects horizontally from a holder located in the aperture.

**Supplementary Video 9. Handle-eat stage of pasta eating in control or optogenetic inhibition trials.** In control trial, the mouse makes coordinated oromanual movements to position and bite the 15mm-angel hair pasta. Optogenetic inhibition of PNs^Emx1^ (4 s - top white bar) of the Rostral Forelimb Orofacial area (RFO) disrupts pasta handling. Mouth orienting to the end of the pasta is interrupted so that eventual biting is directed to the middle of the pasta. Posture is maintained and hand manipulation continues. Optogenetic inhibition of PTs^Fezf2^ (4 s - top white bar) in RFO impairs oromanual manipulation in pasta positioning in the mouth and alters pasta orientation. Optogenetic inhibition of ITs^PlxnD1^ (4 s - top white bar) in RFO results in stiffness of forelimb movement and impairs oromanual coordination to bite/snap the pasta.

## METHODS

### Animals

Adult male and female mice bred onto a C57BL/6J background were used in the experiments. Mice were housed under a 12-h light-dark cycle (7.00 to 19.00 light), with room temperature at 22 °C and humidity at 50%. The experimental procedures were approved by the Institutional Animal Care and Use Committee of Cold Spring Harbor Laboratory (CSHL) and Duke University and performed in accordance with the US National Institutes of Health (NIH) guidelines.

The *Fezf2-CreER*, *Fezf2-Flp*, *PlxnD1-CreER*, *PlxnD1-Flp*, *Sema3E-CreER*, *Tcerg1l-CreER*, *Tbr2-CreER*, and *Tle4-CreER* knock-in mouse driver lines, in which the expression of the inducible Cre recombinase (CreER) or Flp are driven by endogenous promoters, were generated as previously described ^38^. The *Emx1-Cre* knock-in mouse driver line was purchased from Jackson Laboratory (005628). The *Thy1-ChR2* transgenic line 18 (*Thy1-Tg18*) was a gift from Dr. Dinu Florin Albeanu at CSHL. The *Rosa26-loxp-stop-loxp-flpo* (*LSL-Flp*) reporter mice were in-house derived. The *Ai14* (*Rosa26-LSL-tdTomato*), *Ai32* (*Rosa26-LSL-ChR2-eYFP*), *Ai148* (*TIGRE-TRE2-LSL-GCaMP6f-LSL-tTA2*), and *Snap25-LSL-2A-EGFP-D* reporter mice were purchased from Jackson Laboratory (*Ai14*, 007908; *Ai32*, 024109; *Ai148*, 030328; *Snap25-LSL-2A-EGFP-D*, 021879). CreER or Cre driver mice were crossed with *Ai32* or *Ai148* reporter mice for optogenetic stimulation and fiber photometry respectively.

### Viral vectors

The AAV9-Ef1a-DIO-ChR2-eYFP, AAV1-Ef1a-fDIO-GCaMP6f, AAV9-CamKIIa-jGCaMP8m, and AAV9-syn-FLEX-jGCaMP7f-WPRE were purchased from Addgene. The AAV2/8-Ef1a-fDIO-TVA-mCherry, AAV2/8-Ef1a-fDIO-TVA-eGFP, and AAVDJ-DIO-GtACR1-eYFP were produced in house. The AAV8-hSyn-FLEX-TVA-P2A-eGFP-2A-oG and EnVA-dG-Rabies-mCherry were purchased from Salk GT3 Vector Core (La Jolla, California). All viral vectors were aliquoted and stored at -80 °C until use.

### Stereotaxic surgery

Mice, anesthetized with isoflurane (2-5 % at the beginning and 0.8-1.2 % for the rest of the surgical procedure), were positioned in a stereotaxic frame and their body temperature was maintained at 34-37 °C with a heating pad. Lidocaine (2%) was applied subcutaneously to the scalp prior to surgery. Ketoprofen (5 mg/kg) was administered intraperitonially (IP) as an analgesic before and after surgery. A vertical incision was made through the scalp and connective tissue to expose the dorsal surface of the skull. The skin was pushed aside, and the skull surface was cleared using saline. A digital mouse brain atlas, linked to the stereotaxic frame, guided the identification and targeting of different brain areas (Angle Two Stereotaxic System, Leica Biosystems). Coordinates for injections and/or implantations in the RFO were 1.5-1.88 mm anterior from Bregma, 2.25-2.63 mm lateral from midline; aCFA: 0.5 mm anterior from Bregma, 1.5 mm lateral from midline; striatum: 0 mm from Bregma, 3.3 mm lateral from midline, 3 mm ventral from brain surface.

For viral injection, a small burr hole was drilled in the skull and brain surface was exposed. A pulled glass pipette, with a tip of 20-30 μm, containing the viral suspension was lowered into the brain. A 300-400 nl volume was delivered at a rate of 10-30 nl/min using a Picospritzer (General Valve Corp). The pipette remained in place for 5 min, to prevent backflow, prior to retraction. Injections were made at depths of 0.3 and 0.6 mm for *PlxnD1* mice, 0.5 and 0.8 mm for *Fezf2* mice, and 0.3, 0.6, and 0.8 mm for *Emx1* mice. The incision was closed with Tissueglue (3M Vetbond) or 5/0 nylon suture thread (Ethilon Nylon Suture, Ethicon). The mice were kept warm on a heating pad during recovery.

For optogenetic activation, flat optical fibers (diameter 200 μm; NA, 0.22 or 0.39) were implanted in the RFO or striatum. For optogenetic inhibition, flat optical fibers (diameter 400 μm; NA, 0.37) or tapered optical fibers (diameter 400 μm; taper 1 mm; NA, 0.37) were implanted bilaterally in the RFO. For fiber photometry, flat optical fibers (diameter 200 μm; NA, 0.39) were implanted in the right RFO and left aCFA. The tapered optical fibers were implanted at a depth of 550 and 650 μm from the cortical surface in 7 *PlxnD1* and 7 *Fezf2* mice respectively. The flat optical fibers were implanted in the RFO with their tips touching the brain surface. For two *Fezf2* mice used for optogenetic activation, the flat optical fibers were implanted at a depth of 500 μm from the cortical surface. For three *Fezf2* mice used for fiber photometry, the flat optical fibers were implanted at a depth of 400 and 500 μm from the cortical surface in the aCFA and RFO respectively. For drug infusion, two stainless-steel guide cannulae (24-gauge, 62002, RWD Life Science) were implanted bilaterally into the RFO 0.3 mm below the brain surface. To fix the implants to the skull, a silicone adhesive (Kwik-Sil, WPI) was applied to cover the hole, followed by a layer of dental cement (C&B Metabond, Parkell), black instant adhesive (Loctite 426), and dental cement (Ortho-Jet, Lang Dental). A titanium head bar was fixed to the skull near Lambda using dental cement. A plug cannula (62102, RWD Life Science) was inserted into the guide cannula to prevent clogging and reduce the risk of infection.

For thin-skull window preparation, the skull of the right hemisphere was thinned in a 6 mm × 3 mm window preparation (+/- 3 mm AP from Bregma, 3 mm lateral to Bregma) using a micro drill until brain vasculature became visible after saline application. Bregma was then marked in blue. A thin layer of translucent dental cement (C&B Metabond, Parkell) was applied to the thinned skull, followed by nail polish. A titanium head bar was fixed to the skull near Lambda using dental cement (Ortho-Jet, Lang Dental).

For miniscope calcium imaging, the skull covering the right RFO (0.7-2.3 mm anterior from Bregma, 1.1-2.7 mm lateral from the midline) was removed using a micro drill. Cold saline (0.9%) was applied on the skull intermittently during drilling to reduce heat. A 500-600 nl volume of virus was delivered into the RFO (1.5 mm anterior from Bregma, 2.4 mm lateral from the midline) at depths of 0.25, 0.6, and 0.8 mm. A scalpel (#11 blade) was slowly lowered into RFO vertically to a depth of 1.6 mm to create an insertion tract ahead of time. After retracting the scalpel, an integrated prism lens (1 mm diameter, 4.3 mm length; 1000-006614, Inscopix) was disinfected with 100% ethanol, rinsed with 0.9% saline, and slowly inserted into the brain at a depth of 1.4-1.6 mm. To fix the implant to the skull, a silicone adhesive (Kwik-Sil, WPI) was applied to cover the exposed brain, followed by a layer of dental cement (C&B Metabond, Parkell), light-cured dental cement (Maxcem Elite or Variolink Esthetic LC), and black dental cement (Ortho-Jet, Lang Dental). A titanium head bar was fixed to the skull near Lambda using dental cement.

### Tamoxifen induction

Tamoxifen (T5648, Sigma) was dissolved in corn oil (20 mg/ml) by stirring with a magnetic bead at room temperature overnight or by applying a sonication pulse for 60 s, followed by constant rotation overnight at 37 °C. Individual aliquots (1.5 ml each) were stored at 4 °C. For viral injected CreER driver mice, tamoxifen induction was performed via intraperitoneal injections at a dose of 100 mg/kg. The first induction was given one day after the viral injection and subsequent inductions were given once every 2 days for 2-3 times. To drive the reporter gene expression, mice were injected (IP, 100-200 mg/kg) 2-3 times at P21, 28, and/or 35. To label *Tbr2+* neurons born at embryonic day 17 (E17), female and male mice were housed together overnight and females were checked for a vaginal plug between 8-9 am the following morning. Following light isoflurane anesthesia, pregnant females were given oral gavage administration of tamoxifen (dose: 3 mg / 30 g of body weight) at gestational day E17.

### Immunohistochemistry

Adult mice were anaesthetized (using 2.5% Avertin) and intracardially perfused with 25-30 ml PBS followed by 25-30 ml 4% paraformaldehyde (PFA) in 0.1 M PB. After overnight post-fixation at 4 °C, brains were rinsed three times with PBS and sectioned at a thickness of 50-75 μm with a Leica 1000s vibratome. Sections were placed in a blocking solution containing 10% normal goat serum (NGS) and 0.1% Triton-X100 in PBS1X for 1.5 h, then incubated overnight at 4 °C, or room temperature, with primary antibodies diluted in the blocking solution. Sections were rinsed three times (10 min each) in PBS and incubated for 2h at room temperature with corresponding secondary antibodies. Sections were dry-mounted on slides using Fluoromount-G mounting medium (0100-01, SouthernBiotech). Primary antibodies of chicken anti-GFP (1:1,000 or 1:500, Aves, GFP-1020) and rabbit anti-RFP (1:1,000 or 1:500, Rockland Pharmaceuticals, 600-401-379) were used. Alexa Fluor dye-conjugated IgG secondary antibodies (1:500, Molecular Probes, catalog number A11039 for goat anti-chicken 488, A11012 for goat anti-rabbit 594) were used. In some instances, sections were incubated with Neurotrace fluorescent Nissl stain (1:300, Molecular Probes, catalog number N21479) or DAPI (1:1,000, Thermo Scientific, 62248) in secondary antibody. Imaging was performed using a Zeiss Axioimager M2 fluorescence microscope, Zeiss LSM 780 or 710 confocal microscopes, or Zeiss Axio Vert.A1 microscope.

### Retrograde monosynaptic rabies tracing

To map brain-wide monosynaptic inputs onto PTs^Fezf2^ and ITs^PlxnD1^ in the RFO, we first injected the *Fezf2-CreER* or *PlxnD1-CreER* mice with the starter virus of AAV8-hSyn-FLEX-TVA-P2A-eGFP-2A-oG (0.3 μl) in the right RFO. Tamoxifen induction was performed via intraperitoneal injections at a dose of 100 mg/kg, once every 2 days for 3 times (the first induction was one day after the starter virus injection). Three weeks after the AAV injection, mice were injected in the RFO with EnVA-dG-Rabies-mCherry (0.4 μl). Brain tissue was prepared for histologic examination 7-10 days after the rabies virus injection.

Rabies injected brains were imaged either with a Zeiss Axioimager M2 fluorescence microscope or with whole-brain STP tomography. For the wide-field epi-fluorescence imaging, 75-μm coronal sections were obtained across the anteroposterior axis of the brain and every other section was quantitatively analyzed. RFP-labeled (that is, rabies-labeled) input cells were automatically detected, and brain slices were registered to the reference Allen Brain Atlas using Serial Section Registration (http://atlas.brainsmatics.org/a/ssr2021) ^78^. False- and miss-labeled cells were corrected manually. Data are presented as the ratio between the number of RFP-labeled cells in each brain area and the total number of RFP-labeled cells across the entire brain.

### Whole-brain STP tomography

We used the whole-brain STP tomography pipeline previously described^39^. Perfused and post-fixed brains, prepared as described above, were embedded in 4% oxidized agarose in 0.05 M PB, cross-linked in 0.2% sodium borohydrate solution (in 0.05 M sodium borate buffer, pH 9.0-9.5). The entire brain was imaged in coronal sections with a 20× Olympus XLUMPLFLN20XW lens (NA 1.0) on a TissueCyte 1000 microscope (Tissuevision) with a Chameleon Ultrafast-2 Ti:Sapphire laser (Coherent). EGFP/EYFP or tdTomato/mCherry signals were excited at 910 nm or 920 nm, respectively. Whole-brain image sets were acquired as series of 12 (x) × 16 (y) tiles with 1 μm × 1 μm sampling for 230-270 z sections with a 50-μm z-step size. Images were collected by two PMTs (PMT, Hamamatsu, R3896) using a 560 nm dichroic mirror (Chroma, T560LPXR) and band-pass filters (Semrock, FF01-680/SP-25). The image tiles were corrected to remove illumination artifacts along the edges and stitched as a grid sequence. Image processing was completed using Fiji software with linear level adjustments applied only to entire images.

### Axon detection from whole-brain STP data

For axon projection mapping, PN axon signal based on cell-type specific viral expression of EGFP or EYFP was filtered by applying a square root transformation, histogram matching to the original image, and median and Gaussian filtering using Fiji/ImageJ software to maximize signal detection while minimizing background auto-fluorescence ^39^. A normalized subtraction of the autofluorescent background channel was applied and the resulting thresholded images were converted to binary maps. Projections were quantified as the fraction of pixels in each brain structure relative to each whole projection.

### Registration of whole-brain STP image datasets

Registration of brain-wide datasets to the Allen reference Common Coordinate Framework (CCFv3) was performed either by 3D affine registration followed by a 3D B-spline registration using Elastix software, according to established parameters ^39^ or by brainreg software ^79,80^. For axon projection analysis, we registered the CCFv3 to each dataset to report pixels from axon segmentation in each brain structure without warping the imaging channel.

### Axon-projection and monosynaptic-input diagrams from whole-brain imaging data

To generate diagrams of axon projections and monosynaptic inputs for a given driver line, axon- and cell-detection outputs from all individual experiments were compared (sorting the values from high to low) and analyzed side-by-side with low-resolution image stacks (and the CCFv3 registered to the low-resolution dataset for brain area definition) to get a general picture of the injection and high-resolution images for specific brain areas.

### Striatal projection domain analysis

To determine the anatomical and putative functional striatal domains that are targeted by the axons of different cell types in the RFO, we first identified coronal images that match the coronal planes of the data reported in a previous study ^53^. The published striatal images, which have striatal domains annotated, were warped to our corresponding coronal images of the striatum.

### Optogenetic motor mapping

Optogenetic motor mapping techniques were adapted from those previously described (**Fig. 1b**) ^25,26,81^. We briefly anesthetized the mice with isoflurane (2%) to attach a reflective marker on the back of the left hand and to paint their jaw red. Mice were then transferred into a tube, head fixed on a mapping stage, and allowed to fully recover from the anesthesia before stimulation began. The thin-skull window was cleaned with a duster and covered with silicone oil (378399, Sigma-Aldrich). We used a 2D motorized stage (ASI, MS-2000) controlled by MATLAB programs to localize the stimulation at different cortical sites. A 473-nm laser (5-ms pulses, 10 or 50 Hz, 5-20 mW) was used to pseudo-randomly stimulate (100-ms or 500-ms duration) a grid of 128 programmed sites at intervals of 375 µm. A plano-convex lens (focal length (FL) = 250 mm, LA1301-A, Thorlabs) coupled with a SLR photon lens (Voigtlander Nokton, 35 mm FL, f/1.2) was used to collimate the laser beam. The diameter of the laser beam was ∼230 µm (1/e^2^ diameter). A dichroic mirror (Chroma T495lpxr-UF2, round, 2-inch diameter) was used to guide the laser beam to the tissue. Two SLR lenses (the same Nokton 35 mm FL and a Nikkor 105 mm FL, f/2.0, AF), coupled front to front, were used to image the thin-skull window onto the CMOS sensor of a camera (MV1-D1312-40-G2-12, Photonfocus) with a pixel size of 2.67 μm. Bregma was used as the coordinate reference. Each site was stimulated 15-20 times per session. The inter stimulation interval was 2 s. Two cameras (FL3-U3-13E4C-C, FLIR), positioned at the front and the side of the animal, were used to take videorecordings at a frame rate of 100 Hz. The videos were time aligned by TTL signals controlled by the MATLAB programs. The video and TTL-signal states were acquired using workflows in Bonsai software. Four LED light lamps were used for illumination (2 for each camera). After mapping, the thin-skull window was covered with silicone sealant (Kwik-Cast, WPI) for protection and later mapping.

### In vivo optogenetic activation

For head-fixed activation, mice injected with ChR2 virus in the right RFO were prepared and video recorded as described above. For tracking digit movements, two cameras were placed in front of the mice. Two LED light lamps next to the two cameras provided illumination. A fiber coupled laser (5-ms pulses, 5-20 mW; λ = 473 nm) was used to apply stimulation at 10, 20, 30, and 50 Hz and constantly for 0.5 s.

For free-moving activation, mice with optical fibers implanted in the RFO were placed into an acrylic activity box (14 cm × 14 cm × 16.5 cm, L × W × H). A 473-nm laser (5-ms pulses, 5-20 mW) coupled to a rotary joint (RJPFL2, Thorlabs) was used to apply stimulation at 10, 20, and 50 Hz and constantly for 0.5 s. Three cameras (FL3-U3-13S2C-CS, FLIR) were used to take video records at a frame rate of 120 Hz from two sides and the bottom of the activity box. LED light lamps adjacent to each camera provided illumination.

### Pharmacology

The surgery is described in previous sections. A ∼1 mm (medial-lateral) × 1.5 mm (anterior-posterior) craniectomy was made centered around the RFO. The exposed RFO was first covered with 0.9% saline and stimulated to establish baseline movements before drug application. Subsequently, glutamate receptor antagonists, CNQX (4.5 mM) and D-AP5 (2 mM), and GABA_A_ receptor antagonist, gabazine (1 μM) in physiological saline solution were applied to the craniectomy. The compounds were allowed to incubate for 30 min before stimulation resumed, and were replenished (at the same concentration) every ∼10 min throughout the experiment.

### Video analysis for motor mapping and optogenetic activation

Videos of behavior from the motor mapping and head-fixed activation were analyzed either with MATLAB programs or DeepLabCut ^46^. The two cameras were calibrated using the Camera Calibrator App in MATLAB. For hand and jaw tracking in MATLAB, the images were smoothed with a Gaussian low-pass filter (size 9, sigma 1.8). The centroid of the reflective marker on the hand and the tip of the painted jaw were detected by a combination of brightness and hue thresholding, and then tracked by a feature-based tracking algorithm (PointTracker in Computer Vision Toolbox). For DeepLabCut training, 525 images were used from the frontal video record and 700 images were used from the side video record to track the movements of the jaw and hands. For tracking digit movements, an additional set of 750 images acquired by the two front cameras was used in training. Fifteen body parts (the tip and base of the index, middle, and ring fingers of the two hands, the back of the two hands, and the jaw) were labeled in the images. The tracking results were validated manually and errors were corrected accordingly in MATLAB. Trials in which mice made spontaneous movements before stimulation onset (within 0.5 s) were excluded from the analyses, based on either manual examinations or setting threshold (3 × s.d. from the mean) on average speed and acceleration distributions of all trials.

For videos obtained from free-moving activation, the tracking of different body parts was performed using DeepLabCut. The network was trained with 800 images. Eight body parts (left and right eyes, hands, ankles, nose, and tail base) were labeled in the images. The behavioral videos and tracking results were visualized and analyzed in a custom-written MATLAB app. Tracking errors were corrected using the app. For posture analysis, we additionally labeled the back of the mice in the videos manually.

To quantify the hand-to-mouth movement induced by optogenetic activation in the head-fixed animals, we examined the videos from the cortical site that featured the shortest distance between the hand and the nose following the activation (coordinates for 13 *Fezf2* mice: 1.24 ± 0.12 mm anterior from Bregma, 2.39 ± 0.09 mm lateral from the midline; 7 *PlxnD1* mice: 1.66 ± 0.14 mm anterior from Bregma, 2.46 ± 0.11 mm lateral from the midline). Criteria for labeling hand-to-mouth movement were: (1) a forelimb movement that brought the hand to the mouth; (2) a wrist supination; (3) a flexion of the digits. Any intervening grooming movements were not scored as hand-to-mouth movements. We labeled head-to-hand movement by examining the videos from free-moving animals receiving optogenetic stimulation. A head movement that brought the mouth toward the hand contralateral to the stimulation site was defined as a head-to-hand movement.

To compute movement onset time, movement speeds 10 ms before stimulation onset across trials were compared with movement speeds at different time points after stimulation onset by a one-sided paired t-test or one-sided Wilcoxon signed-rank test. The Bonferroni correction was applied to account for multiple comparisons. The earliest post-stimulation time at which a significant increase (p < 0.05) in movement speed was determined was used as movement onset time.

### Angel-hair pasta eating behavior

Mice given ad libitum access to water were food restricted until they reached 80 to 85% of their initial body weight. Food restriction began at least 4 days after surgery. Each day during food restriction, the mice were fed food pellets (0.3-3.5 g of 14 mg Dustless Precision Pellets, F05684, Bio-Serv) to maintain body weight. Most behavioral experiments began after the third day of food restriction, at which time body weights had reached the target level.

Feeding behavior of mice was studied in an automated Mouse Restaurant (**Fig. 2a, Extended Data Fig. 4a**, b). The apparatus has two areas, a dining area (10 cm × 10 cm × 15 cm, dimensions L × W × H) and a waiting area (15 cm × 15 cm × 15 cm), connected by a corridor (24 cm × 4 cm × 15 cm). Food items were placed on a 3D-printed plate mounted on an XZ motorized stage. The plate was moved from the dining area to a food dispenser by two stepper motors (PD42-3-1070, Trinamic Motion Control). The food dispenser, made with two stacked 3D-printed plates, placed a food item onto the table. Each of the two plates, driven by a stepper motor (NEMA-17, 324, Adafruit), could hold 24-food items, such that 48-food items can be provided in the dining area in each session. An acrylic door in the corridor was opened by a servo motor (D625MW, Hitec) to allow access to the dining area from the waiting area. In this way, mice left the waiting area, entered the dining area to eat, and after eating returned to the waiting area where water was accessible from a water port in a corner. Two pairs of infrared (IR) break-beam sensors (2168, Adafruit) installed at each end of the corridor detected the movement direction of the mice. An elevated step fixed between the corridor and the dining area kept food items from being swept out of the dining area. Once mice returned to the waiting area, the door was closed, a new food item was presented and the next trial began. A session ended when all 48-food items had been presented or after 40 minutes, whichever occurred first. The apparatus was controlled by a program running on an Arduino Mega 2560 Rev3 (A000067, Arduino) with three shields (IO sensor shield, DFR0165, DFRobot; LCD and motor shield, 772 and 1438, respectively, Adafruit). An Arduino Uno Rev3 (A000066, Arduino), with three shields (screw shield, DFR0171, DFRobot; LCD and data logging shield, 772 and 1141, respectively, Adafruit), received signals from the IR break-beam sensors to control a laser for optogenetic stimulation and to send TTL signals to recording devices for time alignment.

The mice were pretrained to shuttle between the waiting area and the dining area for one session each day for 2-3 days, were they consumed 30-48, 20-mg pellets (Dustless Precision Pellets, F0163, Bio-Serv). On the day before a pasta-eating session, the mice were familiarized with 0.5 g of angel-hair pasta in their home cage. On the following day, 15-mm pasta pieces were loaded into the food dispenser before the session by inserting them into 3D-printed holders (10 mm × 10 mm × 2 mm, L × W × H, with a 1.5 mm diameter hole in the center, **Extended Data Figs. 4b, 5a**). In the test sessions, the mice consumed 24-48 pieces of 15-mm pasta. During 1-2 sessions, concurrent fiber photometry was obtained. During 6-8 sessions, optogenetic inhibition was applied. At the completion of the 15-mm pasta-eating tests, mice received a training session in which they received 1-mm angel-hair pasta that had been manually placed on the table. Then photometry was obtained over two sessions during which 15-mm and 1-mm pasta lengths, cut using a custom-designed plate, were presented in an alternating order.

### Pasta-bite test

Following the 15-mm pasta-eating sessions, mice used in optogenetic inhibition sessions were given a pasta-bite test. A 20-mm piece of angel-hair pasta was inserted into a metal tube and fixed in place by a screw. The apparatus was located in an aperture (15 mm × 15 mm × 15 mm, L × W × H) made of clear acrylic (**Extended Data Fig. 12a**). A mouse inserted its head into the aperture and bit off pieces of pasta (∼3 mm). One training session was given before the inhibition session. After each trial, mice returned to the waiting area, a new piece of pasta was placed in the holder, and the next trial began. The mice learned to bite the pasta in the first session after which, 1-2 sessions were given with optogenetic inhibition.

### Video recording for pasta eating and data analysis

Three cameras, one on each side of the dining area, video recorded (120 Hz, FL3-U3-13S2C-CS, FLIR) behavior in the dining area. Each camera was fitted with a varifocal lens (T10Z0513CS, Computar). The cameras were time aligned by the TTL signals sent by the Arduino Uno Rev3. The videos and TTL-signal states were acquired using workflows in Bonsai software. Four LED light lamps placed around the dining area provided illumination. Long-pass (590 nm, FGL590S, Thorlabs) and band-pass (435-500 nm, FGB7S, Thorlabs) filters were installed on the light lamps for fiber photometry and miniscope recording respectively. A webcam (C920, Logitech) was installed on a post to monitor the mice from a dorsal perspective.

The cameras were calibrated using the Camera Calibrator App in MATLAB and 14,603 images were pooled to train a deep neural network for tracking using DeepLabCut. Ten body parts (left and right eyes, hands, ankles, nose, tongue, jaw, and tail base) and the pasta (top, center, and bottom) were labeled in the images. The behavioral videos and tracking results were visualized and analyzed in a custom-written MATLAB app.

In the app, we labeled action motifs and sensorimotor events manually through a frame-by-frame analysis (**Extended Data Fig. 4e-i**). Images from all three cameras were displayed for each frame and about 4 million frames were labeled. We identified the start and end frames for the following action motifs: jaw retrieve, tongue lick, sit, left- and right-hand adjustments. The start frame defined movement initiation and the stop frame defined movement completion. A hand-withdraw event was labeled when a mouse raised its hands toward the mouth after chewing. A feeding-end event was labeled when mice lowered their bodies to the floor after food consumption. For saline and muscimol infusion sessions, events in which pasta was dropped were additionally labeled.

In addition to manual labelling, hand-withdraw events and the onsets of chewing were identified with a two-state hidden Markov model (HMM) (https://www.cs.ubc.ca/~murphyk/Software/HMM/hmm.html) using normalized distances of the left- and right-hand to the nose. The model was trained on data from each session with ten random initializations. Only distances from the first bite to the last bite in each trial were used for the training. The model with the largest log likelihood was used to classify the handle-bite and chew phases. A hand-withdraw event was computed as the transition point from a chew phase to a handle-bite phase. Conversely, the onset of chewing was computed as the transition point from a handle-bite phase to a chew phase. We computed the proportions of hand-withdraw events labeled both manually and by the HMM over all hand-withdraw events labeled manually. We found most manually labeled hand-withdraw events were captured by the HMM (88.62 ± 2.53 %, mean ± s.e.m, 6 sessions from 5 mice). In addition, the absolute errors between manually labeled and HMM computed hand-withdraw timestamps were mostly less than 0.1 s (84.13 ± 1.96 %, mean ± s.e.m, 6 sessions from 5 mice).

Action motifs of jaw retrieve, tongue lick, sit, left- and right-hand adjustments were additionally classified using a machine learning pipeline, DeepEthogram ^82^, trained on 509 manually labeled feeding trials. Videos recorded by all three cameras were spatially cropped, concatenated, and downsized frame by frame to a size of 288 × 256 pixels (H × W) for training and inference. TinyMotionNet3D and ResNet3D-34 were used for flow generator and feature extractor respectively. The inference function of the feature extractor module was used for action-motif classification as we found it outperformed the sequence model in classifying hand adjustments. For training and inference, we used one of the following Nvidia GPUs: GeForce 2080Ti or GeForce 4070Ti Super. False-labeled sit was removed manually. False-labeled hand adjustments that were overlapped with bites were removed.

To estimate the time of pasta detection, we first computed the distances from the nose to the top, center, and bottom of the pasta at each onset of pasta retrieval in the control trials. The shortest nose-to-pasta distance at each pasta-retrieval onset was saved. The average shortest distance was used as the pasta-detection distance and computed separately for each mouse. The first time point at which the shortest nose-to-pasta distance drops below the pasta-detection distance was used as pasta-detection time and computed for all trials.

We used Hellinger distance to quantify the similarity between two probability distributions of pasta orientations. For two probability distributions P = (p_1_,…,p_k_) and Q = (q_1_,…q_k_), their Hellinger distance is computed as:

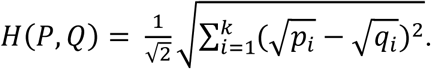

To analyze the phase of hand movements, we first computed the Z-axis movement trajectory using ankle position as a reference. The movement trajectory was band-pass filtered (0.4 - 10 Hz) with forward-backward-zero-phase FIR filters. Hilbert transform was then used on the filtered trajectory to acquire instantaneous phases of the movement. A vector summation was used to obtain the average phase at the time of a bite and the selectivity index of phases:

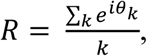

where *θ_k_* is the phase at the time of a bite. The complex phase and amplitude of the resultant R represent the average phase and selectivity index respectively.

### Sound recording and signal analysis

A microphone (AT803, Audio-Technica) on the wall of the dining area picked up the sound of pasta biting. The audio signal from the microphone was amplified (Studio Channel, PreSonus) and digitized at 96,000 Hz by a multifunctional I/O device (PCIe-6323, National Instruments) controlled by MATLAB programs. The TTL signal sent out by the Arduino Uno Rev3 was recorded for time alignment. To detect bite events, the audio signal was band-pass filtered (Butterworth filter, 800-8,000 Hz), rectified, smoothed with a Gaussian window (5 ms), and thresholded (3-5 × s.d. from the mean).

**Muscimol infusion.** After performing 1-2 sessions of 15-mm angel-hair pasta eating, mice were infused with 0.9% saline or muscimol (1 mg/ml, in 0.9% saline) bilaterally into the RFO for two consecutive pasta-eating sessions (saline given for one session, muscimol for the next, or vice versa). Mice were head fixed on a stage and the two hemispheres were infused sequentially after removal of the plug cannula. The injection cannula (28-gauge, 62202, RWD Life Science) connected to a microsyringe (80330, Hamilton) was inserted into the guide cannula to deliver 0.5 or 1 µl of the solution at a rate of 0.1 or 0.2 µl/min by a syringe pump (Legato 130, KD Scientific). After the infusion, the injection cannula was left in place for 5 min to prevent backflow and then retracted, and the plug cannula was reinserted. At the end of the experiments, muscimol diffusion in the brain tissue was determined in two mice by infusing fluorescent muscimol (BODIPY TMR-X Conjugate, 1 mg/ml, dissolved in 50% dimethyl sulfoxide in 0.9% saline; M23400, ThermoFisher Scientific) bilaterally into the RFO (0.5 and 1 µl in the left and right hemispheres respectively), with the same infusion procedure used for the pasta-eating sessions.

### In vivo optogenetic inhibition

The implanted optical fibers were cleaned using alcohol swab sticks and connected to a rotary joint (FRJ_1x2i_FC-2FC, Doric Lenses) with two fiber patch cords (fiber core diameter, 200 μm; RWD Life Science). A fiber coupled laser (5-15 mW; λ = 532 nm) controlled by the Arduino Uno Rev3 was used for the stimulation. For 15-mm angel-hair pasta eating sessions, the laser was turned on for 4 s at mouse entry into the dining area (retrieve-eat stage inhibition, from 0 s to 4 s) or 4 s after entry (handle-eat stage inhibition, from 4 s to 8 s). Thus, the handle-eat stage inhibition targeted oromanual manipulation. We additionally applied 40 Hz stimulation in mice implanted with tapered optical fibers. For sessions of handle-eat stage inhibition, control and inhibition trials in which mice didn’t adopt a sit posture within 4 s were excluded from analysis. For the pasta-bite test, the laser was turned on at entry into the dining area and turned off at the return to the waiting area. Stimulation was given pseudo-randomly for half of the trials in each session.

### Fiber photometry and data analysis

A commercial fiber photometry system (Neurophotometrics) was used to record calcium activity of PTs^Fezf2^ and ITs^PlxnD1^ in the right RFO and left aCFA at 20 Hz. A branching patch cord (fiber core diameter, 200 μm; Doric Lenses) connected the photometry system with the implanted optical fibers. The intensity of the blue light (λ = 470 nm) for GCaMP excitation was adjusted to 20-50 μW at the tip of the patch cord. A violet light (λ = 415 nm, 20-50 μW at the tip) was used to acquire the isosbestic control signal to detect calcium-independent artifacts. Emitted signals were band-pass filtered and focused on the sensor of a CMOS camera. Photometry signals and behavioral events were aligned based on the TTL signals generated by the Arduino Uno Rev3. Mean values of signals from the two regions of interest (ROIs) were calculated and saved by using Bonsai software, and were exported to MATLAB for further analysis.

The recorded photometry signals were processed as previously described ^83,84^. A baseline correction of each signal was made using the adaptive iteratively reweighted Penalized Least Squares (airPLS) algorithm (https://github.com/zmzhang/airPLS) to remove the slope and low frequency fluctuations in the signals. The baseline corrected signals were then standardized (Z-score) on a trial-by-trial basis using the median value and standard deviation of the baseline period (10.6 s, while a mouse is waiting for food delivery). The standardized 415-nm excited isosbestic signal was fitted to the standardized 470-nm excited GCaMP signal using robust linear regression. The standardized isosbestic signal was scaled using parameters of the linear regression and regressed out from the standardized GCaMP signal to obtain calcium dependent signal.

To compute the correlation coefficient between the hand-to-nose distance and GCaMP signal, we used the average of left- and right-hand to nose distances. The hand-to-nose distance was low-pass filtered (5 Hz), shifted forward and backward in time, and downsampled to compute the correlation coefficients of different time lags from -1 s to 1 s. Data in the time window from the first bite to the last bite were used for the correlation analysis.

### In vivo calcium imaging data acquisition and analysis

GCaMP6f/8m fluorescence signals were acquired using a miniature integrated fluorescence microscope system (Inscopix, Palo Alto, CA) through the implanted integrated prism lenses. A motorized commutator (Inscopix) was used to prevent cable entanglement during free-moving recording. Calcium-activity movies were recorded at 20 Hz. The analog gain and LED output power were adjusted based on the dynamic range of the movies. We adjusted the lens focus in the software (IDAS, Inscopix) to bring calcium signals into focus. Recorded calcium signals and behavioral events were aligned based on the TTL signals generated by the Arduino Uno Rev3.

Recorded videos were processed using the IDPS software (Inscopix). We first preprocessed the videos by cropping and spatial downsampling by a factor of 2. The preprocessed videos were band-pass filtered (0.005-0.5 pixel^-1^) and motion corrected. A non-rigid motion correction algorithm (NoRMCorre ^85^) was employed in MATLAB if a global rigid motion correction did not yield acceptable results. Next, we applied the extended constrained nonnegative matrix factorization (CNMF-E), which is optimized for one-photon imaging, to automatically identify the spatial location of cells in the input movie and their associated activity, termed ΔF over noise. CNMF-E identified ROIs were further manually curated in a custom-written MATLAB app to remove ROIs that were not neurons. The ΔF over noise signals were subsequently standardized (Z-score) using the median value and standard deviation of the baseline period (10.6 s, while a mouse is waiting for food delivery).

For clustering the recorded neurons, the average activity of each neuron 0.5 s before and 1.5 s after each actions of step cross, pasta retrieval, hand withdraw, hand adjustment, bite, and chew were computed and normalized to the average activity of a baseline period (-0.5 s to -0.1 s of each action). These average activity were concatenated to form a 2D matrix with rows and columns corresponding to neurons and time points respectively. A PCA analysis was performed to reduce the dimension of the matrix. The first 7 dimensions were retained which account for 91% of the total variance. Spectral clustering analysis was performed on the 7-dimensional activity matrix using MATLAB (*spectralcluster*). The similarity graph for clustering was constructed using the k-nearest neighbour (k-NN) and the cosine metric as the similarity measure for the neuronal activity. A bootstrapping of 100 times, randomly sampling 90% of the data each time, was used to determine the optimal combination of the number of clusters and k-NN. To evaluate the stability of the clustering, we computed the adjusted Rand Index (ARI) and adjusted mutual information (AMI) score for each iteration ^86,87^. Optimal number of clusters and k-NN were subsequently determined based on the maximal mean and the maximal mean adjusted for variance (mean/standard deviation ratio) of ARI and AMI across bootstrapped distributions. The selected parameters enabled reliable clustering as opposed to random assignment of neurons to clusters, as indicated by both the ARI and AMI scores being well above zero^87^.

To determine whether a neuron shows significantly (P < 0.05) increased or decreased activity to an action motif, we performed a paired t-test or Wilcoxon signed-rank test on the data aligned to all onsets (time 0) of that action, which compared the mean Z-scores in two 0.3-s time windows, with one window from -0.4 s to -0.1 s and the other from -0.1 s to 0.2 s.

To decode Z-axis trajectory of the contralateral hand (hand contralateral to the side of recording), the Z-scored calcium activity of simultaneously recorded neurons in each session were used in a linear regression model with ridge regularization (*fitrlinear* in MATLAB). Movement profiles were down sampled to match the sample rate of calcium activity. Explained variance (cvR^2^) was obtained using tenfold cross-validation. A bootstrapping of 1000 times was used to acquire an average cvR^2^ for models including different numbers of recorded neurons as predictors.

A linear logistic regression model with ridge regularization (*fitclinear* in MATLAB) was used to classify the occurrence of bite at different times. The time point at which bite occurs and one time point pre and post to it were labeled as true. A tenfold cross-validation was implemented to obtain precision, recall, and F1 score for the evaluation of model performance. The precision, recall, and F1 score were computed as follows:

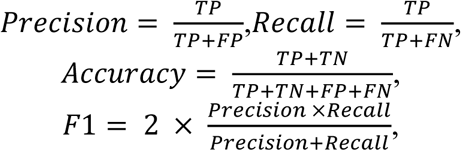

where TP is the number of true positives, FP is the number of false positives, and FN is the number of false negatives. A bootstrapping of 1000 times was used to acquire an average precision, recall, or F1 score for models including different numbers of recorded neurons as predictors.

### Grip and bite strength analysis

Bite strength was measured using an accurate single point load cell system (OEM Style Single Point Load Cells, Omega) ^88^. The system was connected to a custom-built mouth piece with dimensions (H = 3 mm × W = 5 mm × L = 15 mm) based on the incisor morphology of adult C57BL6/J mice. Output signals were amplified (IN-UVI, Omega), digitized via a National Instruments board (PCIe-6323), and fed into a custom MATLAB-based computer interface. A mouse was constrained in a 60-ml plastic tube with an opening on the top to accommodate the implanted cannulae. To prevent the mouse from escaping, a plunger was inserted to loosely confine the mouse. A mouth piece was presented manually and moved slowly at 0.5-1 cm/sec toward the mouth so that the mouse could bite it. Bite strength was measured for 3-4 sessions (120-240 sec per session) for each mouse.

Forelimb grip strength was measured using a custom-designed 3D-printed metal bar (L = 8 cm, diameter = 1.2 mm) attached to an accurate single point load cell system (OEM Style Single Point Load Cells, Omega). The record of the output signal was acquired following a previously described protocol ^89^. In each of 3-4 tests, when a mouse grasped the bar with both hands, its tail was slowly pulled downward with increasing pressure so that the mouse was required to increase its resistance.

### Statistics and data presentation

Significance levels used in the analyses and figures were: *P < 0.05, **P< 0.01, ***P < 0.005, ****P < 0.001, with data presented as mean ± s.e.m., except where otherwise indicated. In the statistical comparisons, data normality was checked with quantile plots and a Shapiro-Wilk normality test in MATLAB. Non-normally distributed data were subsequently compared with non-parametric tests. All statistical tests were two-tailed and adjustments were made for multiple comparisons. No statistical methods were used to predetermine sample size, but our sample sizes are similar to those reported in previous publications ^90,91^.

### Data availability

The data that support the findings of this study are available from the corresponding author upon reasonable request.

### Code availability

Custom-written scripts used in this study are available in a GitHub repository at https://github.com/XuAn-universe/Publication-source-code.

## LIST OF ABBREVIATIONS

ACAd: anterior cingulate area, dorsal part
AId: agranular insular area, dorsal part
APN: anterior pretectal nucleus
CB: cerebellum
CEAc: central amygdalar nucleus, capsular part
CL: central lateral nucleus of the thalamus
CP: caudoputamen
FRP: frontal pole
GPe: globus pallidus, external segment
GPi: globus pallidus, internal segment
GRN: gigantocellular reticular nucleus
GU: gustatory areas
HPF: hippocampal formation
HY: hypothalamus
IRN: intermediate reticular nucleus
MD: mediodorsal nucleus of the thalamus
MdD: medullary reticular nucleus, dorsal part
MDRN: medullary reticular nucleus
MdV: medullary reticular nucleus, ventral part
MOp: primary motor area
MOs: secondary motor area
MRN: midbrain reticular nucleus
OLF: olfactory areas
ORBl: orbital area, lateral part
PAL: pallidum
PARN: parvicellular reticular nucleus
PCN: paracentral nucleus
PF: parafascicular nucleus
PG: pontine gray
PL: prelimbic area
PO: posterior complex of the thalamus
PPN: pedunculopontine nucleus
PSV: principal sensory nucleus of the trigeminal
pyx: pyramidal decussation
RSPagl: retrosplenial area, lateral agranular part
RSPd: retrosplenial area, dorsal part
SC: superior colliculus
SCm: superior colliculus, motor related
SI: substantia innominata
SMT: submedial nucleus of the thalamus
sp: cortical subplate
Spd: spinal cord
SPV: spinal nucleus of the trigeminal
SSp-bfd: primary somatosensory area, barrel field
SSp-ll: primary somatosensory area, lower limb
SSp-m: primary somatosensory area, mouth
SSp-n: primary somatosensory area, nose
SSp-tr: primary somatosensory area, trunk
SSp-ul: primary somatosensory area, upper limb
SSp-un: primary somatosensory area, unassigned
SSs: secondary somatosensory area
STN: subthalamic nucleus
STR: striatum
SUT: supratrigeminal nucleus
V: motor nucleus of trigeminal
VAL: ventral anterior-lateral complex of the thalamus
VII: facial motor nucleus
VISa: anterior area
VISam: anteromedial visual area
VISC: visceral area
VISp: primary visual area
VISpm: posteromedial visual area
VISrl: rostrolateral visual area
VM: ventral medial nucleus of the thalamus
VPM: ventral posteromedial nucleus of the thalamus ZI, zona incerta

